# CD40 is an immune checkpoint regulator that potentiates myocardial inflammation through activation and expansion of CCR2^+^ macrophages and CD8 T-cells

**DOI:** 10.1101/2024.03.14.584418

**Authors:** Jesus Jimenez, Junedh Amrute, Pan Ma, Xiaoran Wang, Raymond Dai, Kory J. Lavine

**Affiliations:** Center for Cardiovascular Research, Department of Medicine, Cardiovascular Division, Washington University School of Medicine, St. Louis, MO, USA; Cardio-Oncology Center of Excellence, Department of Medicine, Cardiovascular Division, Washington University School of Medicine, St. Louis, MO, USA; Washington University, St. Louis, MO, USA; Department of Pathology and Immunology, Washington University School of Medicine, St. Louis, MO, USA; Department of Developmental Biology, Washington University School of Medicine, St. Louis, MO, USA

**Keywords:** cardio-oncology, CD40, immune checkpoint inhibitor, macrophages, cardiac immunology, interferon gamma, CD8 T-cells

## Abstract

Novel immune checkpoint therapeutics including CD40 agonists have tremendous promise to elicit antitumor responses in patients resistant to current therapies. Conventional immune checkpoint inhibitors (PD-1/PD-L1, CTLA-4 antagonists) are associated with serious adverse cardiac events including life-threatening myocarditis. However, little is known regarding the potential for CD40 agonists to trigger myocardial inflammation or myocarditis. Here, we leveraged genetic mouse models, single cell sequencing, and cell depletion studies to demonstrate that an anti-CD40 agonist antibody reshapes the cardiac immune landscape through activation of CCR2^+^ macrophages and subsequent recruitment of effector memory CD8 T-cells. We identify a positive feedback loop between CCR2^+^ macrophages and CD8 T-cells driven by IL12b, TNF, and IFN-γ signaling that promotes myocardial inflammation and show that prior exposure to CD40 agonists sensitizes the heart to secondary insults and accelerates LV remodeling. Collectively, these findings highlight the potential for CD40 agonists to promote myocardial inflammation and potentiate heart failure pathogenesis.

## Introduction

Cardiovascular disease is the predominant risk factor for morbidity and mortality among cancer survivors^1, 2^. Cancer treatments directly result in cardiovascular toxicity and may exacerbate comorbid diseases such as hypertension and diabetes. Among cancer treatments contributing to survivorship, immunotherapies including immune checkpoint inhibitors (ICIs) are now widely utilized due to their groundbreaking efficacy.

ICIs promote tumor cell death by activating T-cell responses through modulation of ligand-receptor interactions between T-cells and antigen presenting cells such as macrophages and dendritic cells, or cancer cells. Conventional PD-1 (programmed cell death 1), PD-L1 (programmed death ligand-1), and CTLA-4 (cytotoxic T-lymphocyte-associated protein 4) antagonistic antibodies block an inhibitory signal restraining T-cell activation. These agents impose substantial cardiac risk ranging from arrhythmias and reduced cardiac function to fulminant myocarditis, which affects at least 1% of patients and is associated with high mortality^3, 4^. Current treatments for ICI myocarditis are lacking and limited to high dose steroids and anecdotal use of intravenous abatacept/belatacept, immunoglobulin, plasmapheresis, and rituximab^5, 6^.

Macrophages represent the predominate antigen presenting cell. Under steady state conditions, the heart contains two major subsets of macrophages with differing origins and functions: tissue resident macrophages and monocyte-derived macrophages. Among these populations, monocyte-derived macrophages express C-C chemokine receptor 2 (CCR2) and contribute to myocardial inflammation and heart failure pathogenesis^7, 8^. CCR2^+^ macrophages activate a robust inflammatory response through the generation of proinflammatory cytokines and chemokines, recruitment of innate immune cells, and activation of T-cells. ICI myocarditis is characterized by infiltration of CCR2^+^ macrophages and activated CD8 T-cells into the heart. Each of these cell types is causatively linked to disease pathogenesis^9–11^.

Given the success of conventional ICIs, considerable effort has focused on identifying additional immune checkpoint pathways that could serve as cancer therapeutic targets such as CD40, OX40, and LAG-3. Here, we focused on the CD40 pathway, which represents an activating signal between T-cells and antigen presenting cells that elicits antitumor responses, even in patients resistant to established treatments including conventional immune checkpoint blockade^12, 13^. These results have led to the development of CD40 activating (agonist) antibodies. The potential of acute CD40 signaling activation to drive myocardial inflammation or myocarditis is unclear.

Here, we investigated whether activation of CD40 signaling reshapes the cardiac immune landscape and triggers myocardial inflammation. We utilized an anti-CD40 antibody that stimulates receptor activation mimicking the clinical therapy under investigation. Using genetically engineered mouse models in conjunction with single cell RNA sequencing, molecular pathology, and cell depletion studies, we defined the inflammatory response provoked by CD40 activation and underlying mechanistic determinants. We identified expansion of inflammatory CCR2^+^ macrophages and CD8 T-cells with an effector memory phenotype within the myocardium following treatment with the anti-CD40 agonist antibody. There was increased myocardial expression of numerous inflammatory cytokines and chemokines. We demonstrated that anti-CD40 agonist antibody directly activates CCR2^+^ macrophages, which triggered the recruitment and activation of IFN-γ producing effector memory CD8 T-cells through IL-12b and TNF signaling. We identified a positive feedback loop between CD8 T-cells and CCR2^+^ macrophages driven by IFN-γ signaling that was required for myocardial inflammation. Finally, we showed that prior exposure to anti-CD40 agonist antibody sensitizes the heart to secondary hypertensive and ischemic insults and accelerates LV remodeling. Collectively, these findings establish the potential for CD40 signaling to trigger myocardial inflammation and accelerate heart pathogenesis. By defining the underlying cellular and signaling mechanisms, we identify putative targets to suppress myocardial inflammation provoked by CD40 agonists.

## Results

### CD40 agonist treatment triggers expansion of immune cells in the heart

To investigate the effect of anti-CD40 agonist antibody treatment in the murine heart, wildtype (WT) C57BL/6J mice received serial intraperitoneal (i.p.) injections with anti-CD40 (100 µg every 72 hrs) for 7 days (**Fig. 1a**). Hematoxylin and eosin (H&E) staining of formalin-fixed paraffin-embedded (FFPE) myocardial sections demonstrated expansion of immune cells (**Fig. 1b**). Since CD40 agonism activates antigen presenting cells, we investigated the composition of the immune cells starting with macrophages, which represent the predominate antigen presenting cell within the heart. CD64 immunostaining revealed increased macrophage abundance (**Fig. 1c**). CD8 immunostaining also revealed increased T-cell abundance (**Fig. 1d**) after 7 days of CD40 agonist treatment. Despite evidence of myocardial inflammation, there was no significant change in left ventricular (LV) chamber size or LV systolic function in mice treated with anti-CD40 agonist antibody (**Extended Data Fig. 1a-c**).

**Fig. 1.**
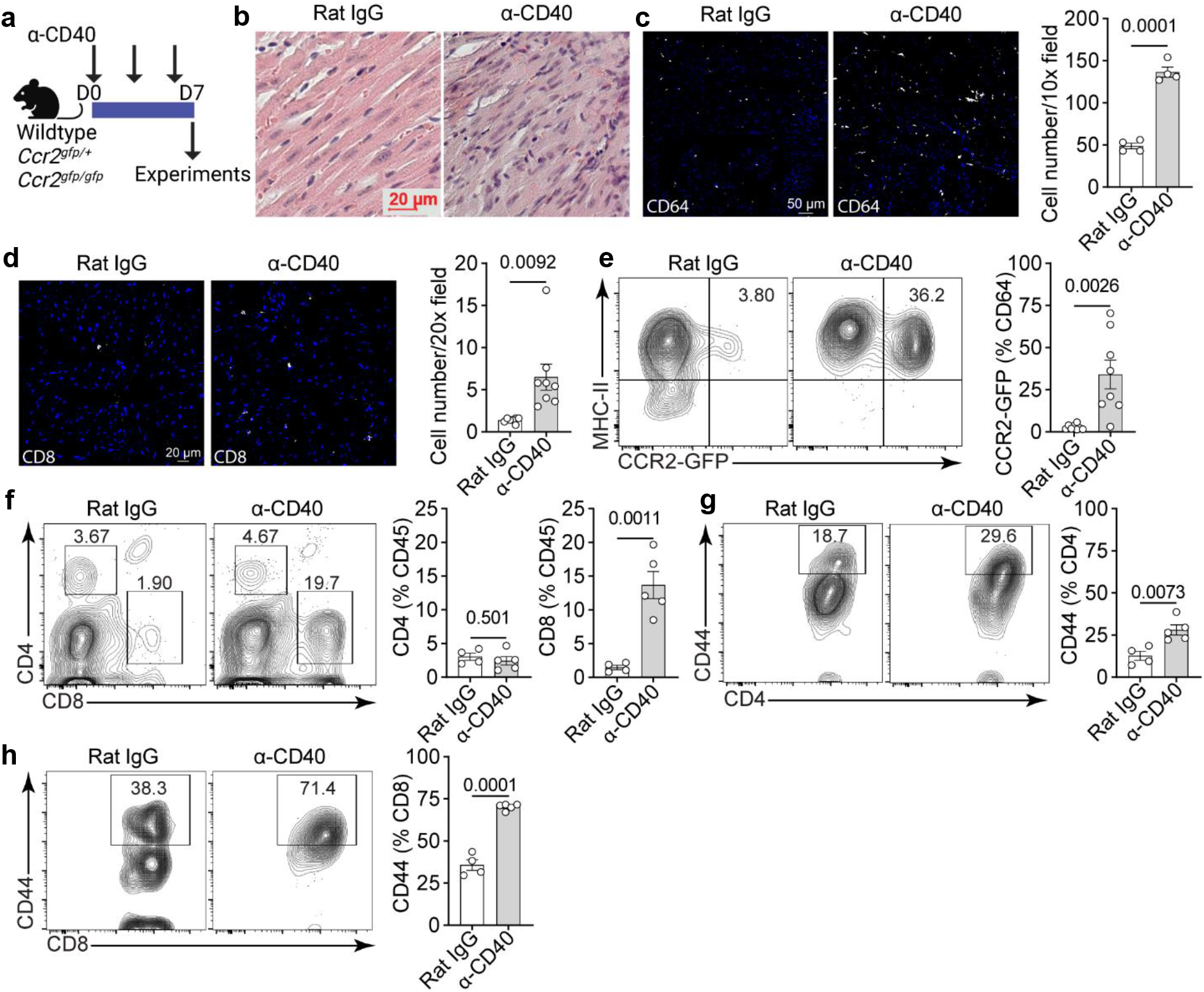
Expansion of CCR2^+^ macrophages and effector memory T-cells in anti-CD40 agonist antibody treated hearts. **a,** Experimental model of mice receiving α-CD40 agonist treatment or Rat IgG isotype antibodies. **b,** Representative hematoxylin and eosin (H&E) staining showing increased immune cells in the wildtype mouse hearts comparing Rat IgG to α-CD40. Scale bar 20 µm. **c,** Representative CD64 macrophage immunofluorescent staining (white) in wild-type mouse hearts, and quantification of CD64^+^ cells per 10x field. Comparison between Rat IgG (n = 4) and α-CD40 (n = 4). Scale bar 50 µm. **d,** Representative CD8 immunofluorescent staining (white) in wild-type mouse hearts and quantification of CD8^+^ cells per 20x field. Comparison between Rat IgG (n = 7) and α-CD40 (n = 8). Data collected from 2 independent experiments. Scale bar 20 µm. **e,** Quantification of CCR2-GFP^+^ macrophages in the heart by flow cytometry comparing Rat IgG (n = 8) and α-CD40 (n = 8). Data collected from 2 independent experiments. **f,** Quantification of CD4^+^ and CD8^+^, **g,** CD4^+^CD44^+^ and **h,** CD8^+^CD44^+^ T-cells in the heart by flow cytometry comparing Rat IgG (n = 4) and α-CD40 (n = 5). Two-tailed unpaired Student’s *t*-test performed for all statistical analysis. Error bars indicate means ± s.e.m.

To further differentiate which macrophage subsets expand following anti-CD40 agonist antibody treatment, hearts were processed to generate single cell suspensions and prepared for flow cytometry analysis (**Extended Data Fig. 2**). *Ccr2^gfp/+^*reporter mice were used as they are a robust tool to identify CCR2^+^ macrophages based on GFP florescence. We found a significant increase in GFP-labeled CCR2^+^ macrophages, consistent with expansion of pro-inflammatory macrophages within the heart (**Fig. 1e**). To determine if classic dendritic cells also expanded within the heart, *Zbtb46^gfp/+^* mice^14^ were treated with anti-CD40 agonist antibody for 7 days. Flow cytometry analysis demonstrated minimal increase in GFP-labeled dendritic cells (**Extended Data Fig. 3a**). There was not a significant difference in neutrophils or B cells (**Extended Data Fig. 3b-c**). We performed additional flow cytometry studies to further characterize expanded T-cells within the heart following anti-CD40 agonist antibody treatment. We found a significant increase in CD8 T-cells with an effector memory phenotype (**Fig. 1f-g**). There was also a significant increase in CD4 effector memory T-cells (**Fig. 1f**, **1h**). These findings support that anti-CD40 agonist antibody treatment results in expansion of CCR2^+^ macrophages and effector memory phenotype CD4 and CD8 T-cells in the heart.

### Dynamics of CCR2^+^ macrophage and T-cell expansion and dependence on monocyte recruitment

To determine if expansion of CCR2^+^ macrophages and T-cells within the heart requires monocyte recruitment, we utilized a tamoxifen-inducible monocyte lineage tracing strategy and examined CCR2 deficient mice. We have previously demonstrated that administration of tamoxifen to *Ccr2^ertCre^Rosa26^tdTomato^*mice results in labeling of over 80% of peripheral monocytes and without labeling macrophages resident within the heart^15^. To delineate whether infiltrating monocytes contribute to the expansion of CCR2^+^ macrophages observed following anti-CD40 agonist antibody treatment, we administered tamoxifen (60mg/kg every 3 days) to *Ccr2^ertCre^Rosa26^tdTomato^* mice. Mice were concurrently treated with either isotype control or anti-CD40 agonist antibody (**Fig. 2a**). We observed a significant increase in tdTomato-labeled CCR2^+^ macrophages, indicating that monocyte recruitment contributes to CD40 agonist driven expansion of CCR2^+^ macrophages (**Fig. 2b**). To determine if monocyte recruitment is required for expansion of CCR2^+^ macrophages, we examined control (*Ccr2^gfp/+^*) and CCR2 deficient (*Ccr2^gfp/gfp^*) mice, which allow visualization of CCR2^+^ cells in the absence of CCR2 protein expression. Following treatment with anti-CD40 agonist antibody for 7 days, flow analysis revealed a significant increase in CCR2^+^ macrophages in anti-CD40-treated control and *Ccr2^gfp/gfp^* hearts compared to isotype control treated hearts. There was not a significant difference of CCR2^+^ macrophages between anti-CD40-treated control and *Ccr2^gfp/gfp^* hearts (**Fig. 2c**). Similarly, control and *Ccr2^gfp/gfp^* hearts displayed a similar increase in CD4 and CD8 effector memory T-cells following anti-CD40 agonist antibody treatment (**Fig. 2d-h**).

**Fig. 2.**
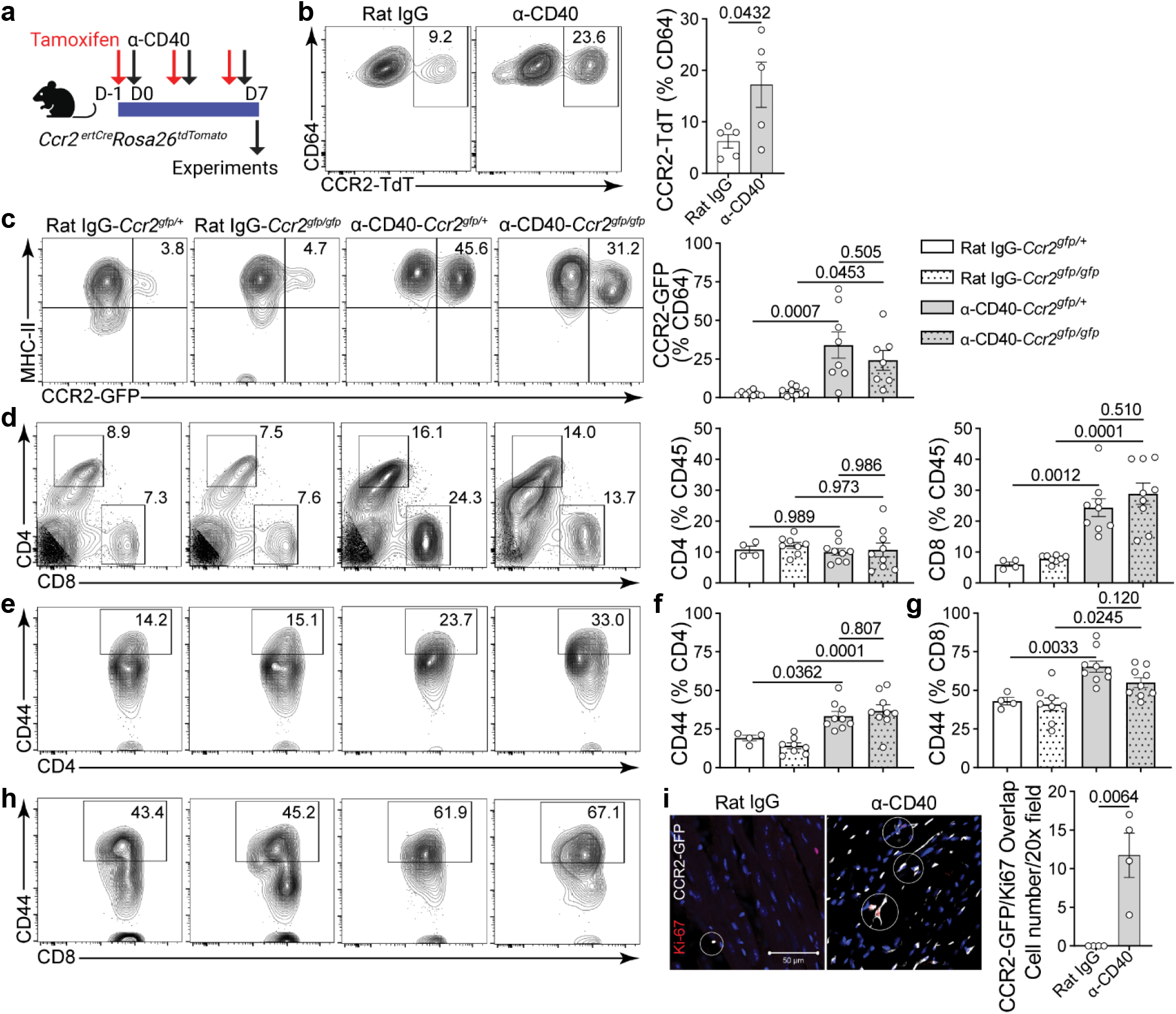
Dynamics of CCR2^+^ macrophage and T-cell expansion and dependence on *monocyte recruitment.* **a,** Experimental model with mice receiving tamoxifen to label CCR2 macrophages and concurrent treatment with α-CD40 agonist or Rat IgG isotype. **b,** Quantification of CCR2-TdT^+^ macrophages in the heart by flow cytometry between Rat IgG (n = 5) and α-CD40 (n = 5), two-tailed unpaired Student’s *t*-test. **c,** Quantification of CCR2-GFP^+^ macrophages in the heart by flow cytometry in control (*Ccr2^gfp/+^*) and CCR2 deficient (*Ccr2^gfp/gfp^*) mice. Comparisons are between Rat IgG-*Ccr2^gfp/+^* (n = 8), Rat IgG-*Ccr2^gfp/gfp^* (n = 8), α-CD40-*Ccr2^gfp/+^* (n = 8), and α-CD40-*Ccr2^gfp/gfp^* (n = 7), one-way ANOVA with Sidak correction. Data collected from 2 independent experiments **d,** Quantification of CD4 and CD8 T-cells, **e, f,** CD4^+^CD44^+^ T-cells, and **g, h,** CD8^+^CD44^+^ T-cells in the heart by flow cytometry comparing Rat IgG-*Ccr2^gfp/+^* (n = 4), Rat IgG-*Ccr2^gfp/gfp^* (n = 8), α-CD40-*Ccr2^gfp/+^* (n = 9), and α-CD40-*Ccr2^gfp/gfp^* (n = 9), one-way ANOVA with Sidak correction. Data collected from 2 independent experiments. **i,** Ki67 (red) and CCR2-GFP (white) immunofluorescent staining in wildtype mice comparing Rat IgG (n = 4) and α-CD40 (n = 4), two-tailed unpaired Student’s *t*-test. White circles highlight overlap of Ki67 and CCR2-GFP. Scale bar 50 µm. Error bars indicate means ± s.e.m.

We performed immunostaining using Ki67 to assess whether macrophage proliferation also contributes to CCR2^+^ macrophage expansion^8, 16^. Indeed, there was increased Ki67 immunostaining that overlapped with CCR2^+^ cells in heart sections of mice treated with anti-CD40 agonist antibody, consistent with CCR2^+^ macrophage cell proliferation (**Fig. 2i**). These findings demonstrate that CCR2^+^ macrophages expand via both monocyte recruitment and cell proliferation in response to anti-CD40 agonist antibody treatment. However, monocyte recruitment is not necessary for CCR2^+^ macrophage or T-cell expansion.

### CD40 activation promotes myocardial inflammatory cytokine and chemokine production

To define the cytokine and chemokine signature driving expansion of CCR2^+^ macrophages and effector memory T-cells within the heart, we performed RT-qPCR from bulk cardiac tissue of wildtype mice treated with anti-CD40 agonist antibody. We found a significant increase in multiple inflammatory cytokines including *Ifng*, *Il12b*, and *Tnf* as well as inflammatory chemokines including *Ccl5*, *Cxcl9*, and *Cxcl10*. Many of these effectors are regulated by IFN-γ and function to coordinate antigen presenting cell and T-cell activation^17–19^ (**Fig. 3a**). We have previously demonstrated that in ICI myocarditis CCR2^+^ macrophages generate CXCL9^10^. To determine if CCR2^+^ macrophages also express *Cxcl9* in response to anti-CD40 agonist antibody treatment, we performed *in situ* RNA hybridization with RNA probes for *Ccr2* and *Cxcl9*. We observed increased expression of *Cxcl9* that co-localized with CCR2^+^ macrophages (**Fig. 3b**). These findings demonstrate that following anti-CD40 agonist antibody treatment, there is marked upregulation of inflammatory cytokines and chemokines within the heart and suggests the possibility of inflammatory crosstalk between CCR2^+^ macrophages and T-cells.

**Fig. 3.**
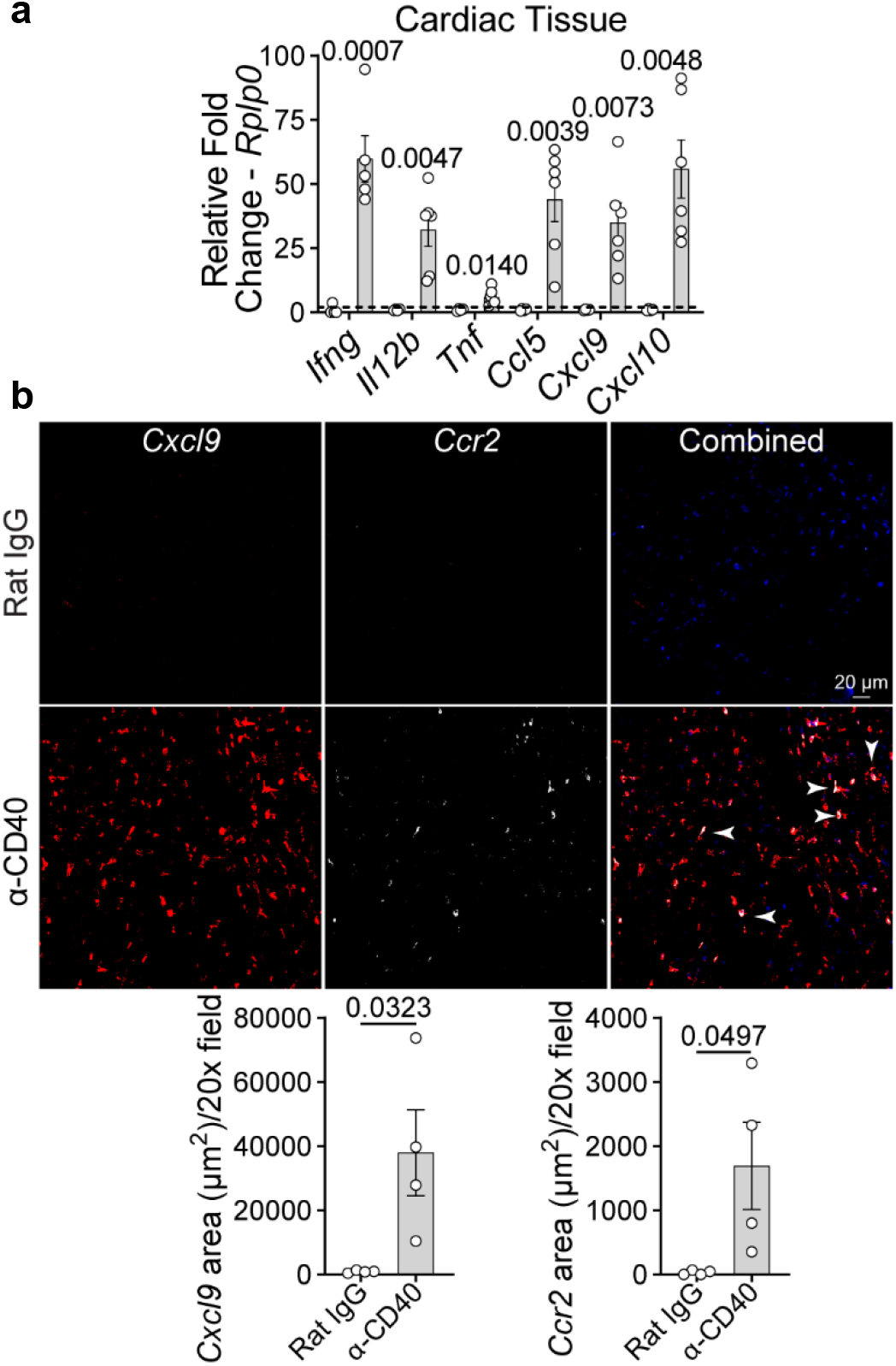
CD40 activation promotes myocardial inflammatory cytokine and chemokine *production.* **a,** *Ifng*, *Il12b*, *Tnf*, *Ccl5*, *Cxcl9*, and *Cxcl10* mRNA expression measured by RT-qPCR from wildtype bulk cardiac tissue. Comparison between Rat IgG (n = 4) and α-CD40 (n = 5-6) for each individual gene. **b,** Representative images of *Cxcl9* (red) and *Ccr2* (white) expression detected in Rat IgG (n = 4) and α-CD40 (n = 4) of wildtype mouse hearts by RNA *in situ* hybridization. Co-expression noted with white arrowheads, scale bar 20 µm. Quantification of *Cxcl9* and *Ccr2* fluorescent area per 20x field. Two-tailed unpaired Student’s *t*-test performed for all statistical analysis. Error bars indicate means ± s.e.m.

### CD40 signaling drives transcriptional shifts in mononuclear phagocytes

To more deeply explore shifts in the cardiac immune landscape triggered by anti-CD40 agonist antibody treatment, we performed cellular indexing of transcriptomes and epitopes sequencing (CITE-seq) on CD45^+^ cells isolated from WT hearts following 7 days of isotype or anti-CD40 agonist antibody treatment (**Extended Data Fig. 4a-e**). Unsupervised clustering identified 4 major immune cell populations including mononuclear phagocytes (monocytes, macrophages, dendritic cells), T-cells and natural killer (NK)-cells, B cells, and neutrophils (**Fig. 4a**). Subclustering of mononuclear phagocytes identified 9 transcriptional cell states including 5 groups of macrophages, 3 groups of monocytes, and a group of classical dendritic cells (**Fig. 4b**). Differential expression analysis identified the following specific marker genes: Mac1 (*Ccl8*, *Cxcl9, Ccl7*), Mac2 (*F13a1*, *Cd163, Lyve1*), Mac3 (*Ly6a*, *Ccr2*, *Mmp14*), Mac4 (*Cxcl10, Cxcl2, Ccl4*), Mac5 (*Ccl5, Cxcl9, Il12b*), classical monocytes 1 (*Plac8*, *Chil3*, *Ly6c2*), classical monocytes 2 (*Fn1*, *Chil3*, *Lsp1*), non-classical monocytes (*Ear2*, *Nr41*, *Ace*), and classic dendritic cells (*Cd209*, *Cst3*, *Lsp3*) (**Fig. 4c**).

**Fig. 4.**
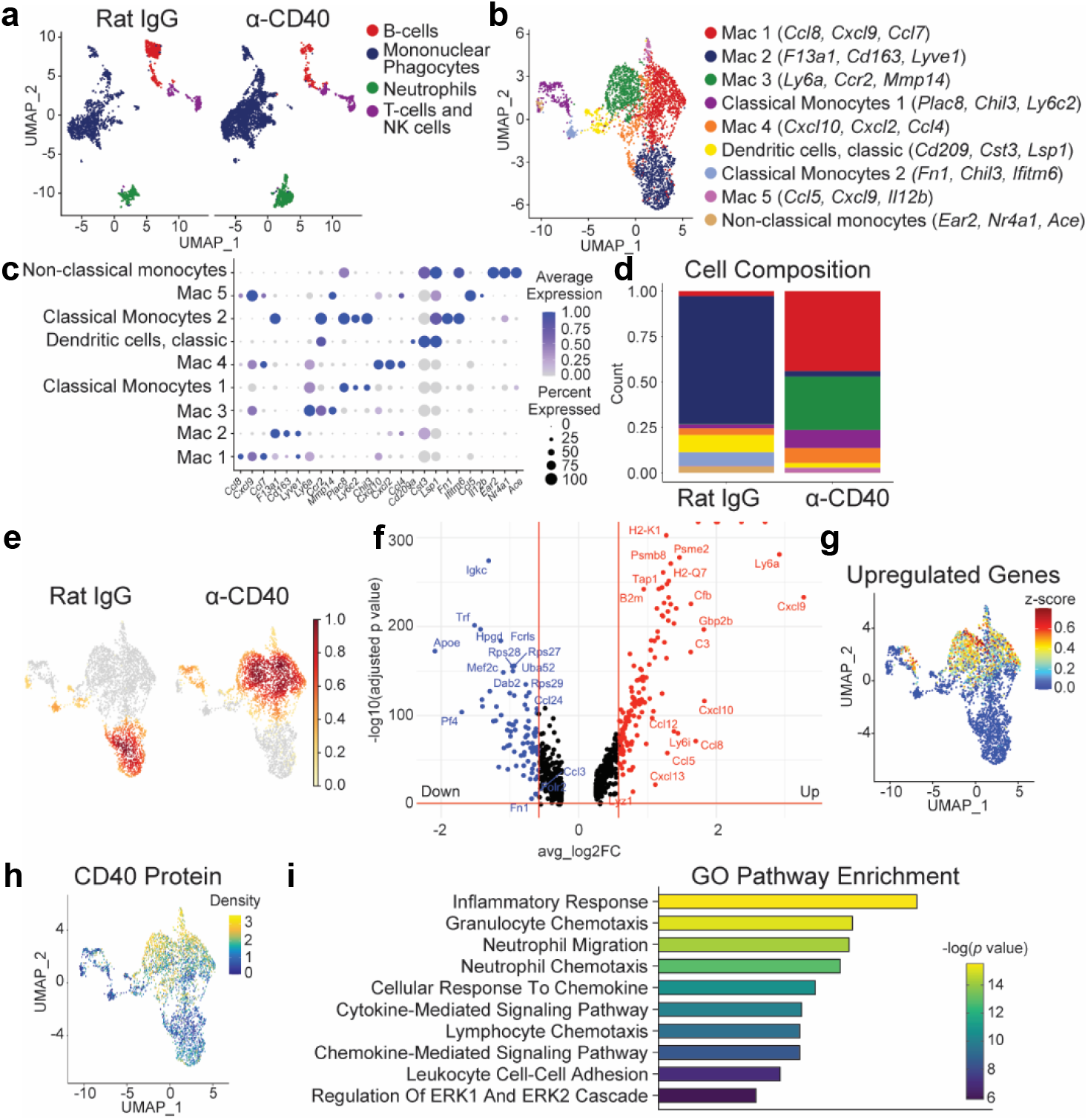
CD40 signaling drives transcriptional shifts in mononuclear phagocytes. **a,** UMAP clustering from a total of 8 hearts treated with Rat IgG (n = 4) or α-CD40 (n = 4) showing 4 major cell types split by experimental group. **b,** UMAP subclustering of mononuclear phagocytes highlighting 9 transcriptionally distinct subclusters. **c,** Dot plots of differentially expressed genes in each mononuclear phagocyte subcluster. **d,** Cell composition proportion and **e,** Gaussian kernel density plots of mononuclear phagocyte nuclei comparing Rat IgG and α-CD40. **f,** Volcano plot of differentially expressed genes comparing Rat IgG to α-CD40 agonist treated hearts. Analyzed using Wilcoxon rank-sum test using R package Seurat (v4). **g,** Z-score feature plot of top upregulated genes in mononuclear phagocytes. **h,** Density plot showing expression CD40 protein in UMAP embedding in mononuclear phagocytes. **i,** Gene Ontology (GO) pathway enrichment analysis of upregulated genes in mononuclear phagocytic cells displaying the top 10 enriched pathways. Genes used in the analysis were selected from Seurat differential expression with an adjusted *p* < 0.05 and absolute (log_2_FC > 1.0).

We observed striking shifts in mononuclear cell states between treatment groups (**Fig. 4d-e**). Mac2 (*F13a1*, *Cd163, Lyve1*), which expresses canonical markers of tissue resident macrophages^20, 21^, were markedly reduced in hearts treated with anti-CD40 agonist antibody compared to isotype control treated hearts. We also detected reductions in classical monocytes 2, non-classical monocytes, and dendritic cells in anti-CD40 agonist antibody treated hearts. In contrast, hearts treated with anti-CD40 agonist antibody demonstrated robust enrichment of Mac1, Mac3, Mac4, Mac5, and classical monocytes 1 compared to isotype control treated hearts. Differential gene expression analysis of mononuclear phagocytes isolated from isotype control treated versus anti-CD40 agonist treated hearts revealed increased expression of 225 genes that included numerous pro-inflammatory mediators (**Fig. 4f**). Many of the top differentially expressed genes that were expressed in macrophage (Mac1, Mac3, Mac4, Mac5) and monocyte (classical monocytes 1) clusters were enriched with CD40 protein expression following anti-CD40 agonist antibody treatment (**Fig. 4g-h**). Gene Ontology (GO) pathway enrichment analysis using genes upregulated in anti-CD40 agonist treated mononuclear phagocytes highlighted that inflammatory response, cellular response to chemokine, and cytokine- and chemokine-mediated signaling pathway were among the topmost affected pathways (**Fig. 4i**). Density plots revealed that *Cxcl9, Il12b,* and *Tnf* were expressed in CD40^+^ monocytes and macrophages isolated from hearts treated with anti-CD40 agonist antibody (**Figure 4H**, **Extended Data Fig. 5a-c**).

### CD40 signaling triggers transcriptional shifts in T-cells and NK-cells

To further evaluate the cardiac immune landscape, we performed additional subclustering of T-cells and NK-cells and identified 6 subpopulations based on transcriptomic and cell surface protein features including effector memory CD8 T-cells, 4 groups of NK-cells, and a naïve T-cell cluster (**Fig. 5a-b**)^22^. Density plots of CD3, CD8, and CD4 proteins confirmed enriched expression in the T-cell subpopulations (**Fig. 5c-e**). Cell composition and gaussian kernel density plots demonstrated a marked shift from naïve T-cells to CD8 T-cells and from NK-cell state 1 to NK-cell states 2, 3, and 4 (**Fig. 5f-g**). Differential gene expression analysis identified multiple inflammatory mediators that were upregulated in T-cells and NK-cells from hearts treated with anti-CD40 agonist antibody compared to isotype control treated hearts (**Fig. 5h**). Z-score feature plot of the top upregulated genes indicated expression in most populations with the exception of naïve T-cells (**Fig. 5i**). GO pathway enrichment analysis using genes upregulated in anti-CD40 agonist treated T-cells and NK-cells highlighted that cellular response to cytokine stimulus including type II interferon, and alpha-beta T-cell activation were among the topmost affected pathways (**Fig. 5j**). Among the upregulated genes and pathways, *Ifng* showed a robust increase in CD8^+^ T-cells and NK-cells from hearts exposed to anti-CD40 agonist antibody (**Fig. 5k-l**).

**Fig. 5.**
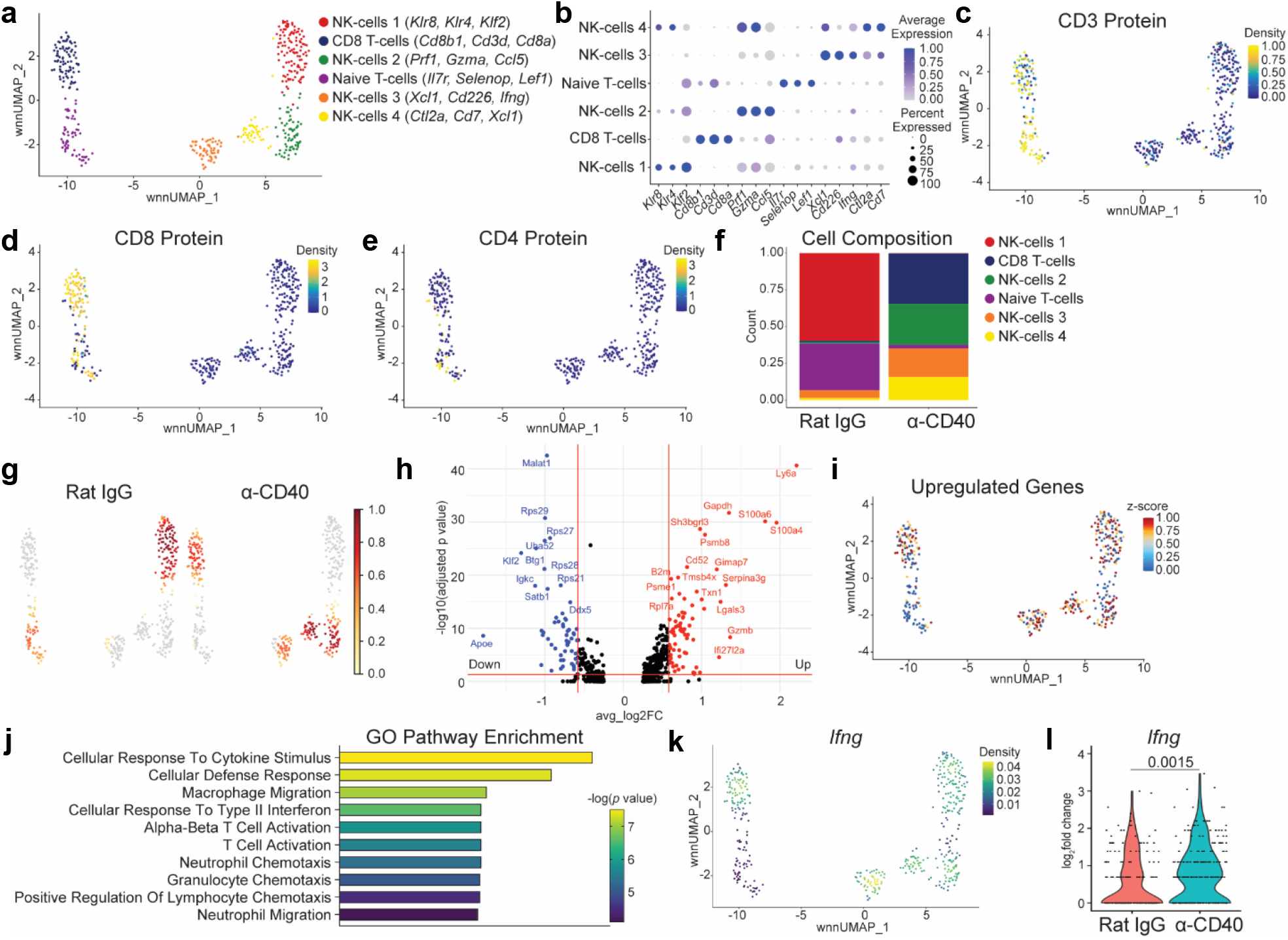
CD40 signaling triggers transcriptional shifts in T-cells and NK-cells. **a,** Weighted nearest neighbor (wnn) UMAP subclustering of T-cells and NK-cells split by experimental group highlighting 6 transcriptionally distinct subclusters. **b,** Dot plots of differentially expressed genes in each T-cell and NK-cell subcluster. **c,** Density plot showing expression of CD3, **d,** CD8, and **e,** CD4 protein in wnnUMAP embedding in T-cells and NK-cells. **f,** Cell composition proportion and **g,** Gaussian kernel density plots of T-cell and NK-cell nuclei comparing Rat IgG and α-CD40. **f,** Volcano plot of differentially expressed genes comparing Rat IgG to α-CD40 agonist treated hearts. Analyzed using Wilcoxon rank-sum test using R package Seurat (v4). **i,** Z-score feature plot of top upregulated genes in T-cells and NK-cells. **j,** Gene Ontology (GO) pathway enrichment analysis of upregulated genes in T-cells and NK-cells displaying the top 10 enriched pathways. **k,** Density plot showing expression of *Ifng* in wnnUMAP embedding in T-cells and NK-cells. **l,** Violin plot of *Ifng* expression comparing Rat IgG and α-CD40, two-tailed unpaired Student’s *t*-test. Genes used in the analysis were selected from Seurat differential expression with an adjusted *p* < 0.05 and absolute (log_2_FC > 1.0).

### CD40 signaling in CCR2^+^ macrophages drives expansion and activation of CCR2^+^ macrophages and CD8 T-cells

Next, we sought to determine whether CD40 signaling in CCR2^+^ macrophages is responsible for CCR2^+^ macrophage expansion and CD8 T-cell recruitment, activation, and IFN-γ production. To examine this possibility, we conditionally deleted CD40 from CCR2^+^ macrophages by generating *Cd40^flox/flox^Ccr2^ertCre^*mice (*Cd40^flox/flox^* generously shared by Dr. Esther Lutgens^23^). Mice received tamoxifen (i.p. injections 40 mg/kg tamoxifen daily for 5 days followed by tamoxifen food pellets) for two weeks followed by anti-CD40 agonist antibody treatment for 7 days (**Fig. 6a**). Using flow cytometry analysis, we observed complete attenuation of CCR2^+^ macrophage expansion and CD8 effector memory T-cells in *Cd40^flox/flox^Ccr2^ertCre^* hearts compared to controls treated with anti-CD40 agonist antibodies (**Fig. 6b-f**). Consistent with these findings, upregulation of *Ifng* and *Cxcl9* expression observed in control hearts treated with anti-CD40 agonist antibody relative to isotype treated hearts was abrogated in *Cd40^flox/flox^Ccr2^ertCre^*hearts (**Fig. 6g-h**). Since CD40 signaling plays an essential role in the humoral immune response through B cell activation^24^, we also sought to determine the effect of depleting B cells on anti-CD40 agonist antibody treated mice. B-cell depletion had no effect on expansion of cardiac CCR2^+^ macrophages in response to anti-CD40 agonist antibody treatment (**Extended Data Fig. 6a-b**). These findings reveal that the anti-CD40 agonist antibody exerts its inflammatory effects by directly signaling to CCR2^+^ macrophages.

**Fig. 6.**
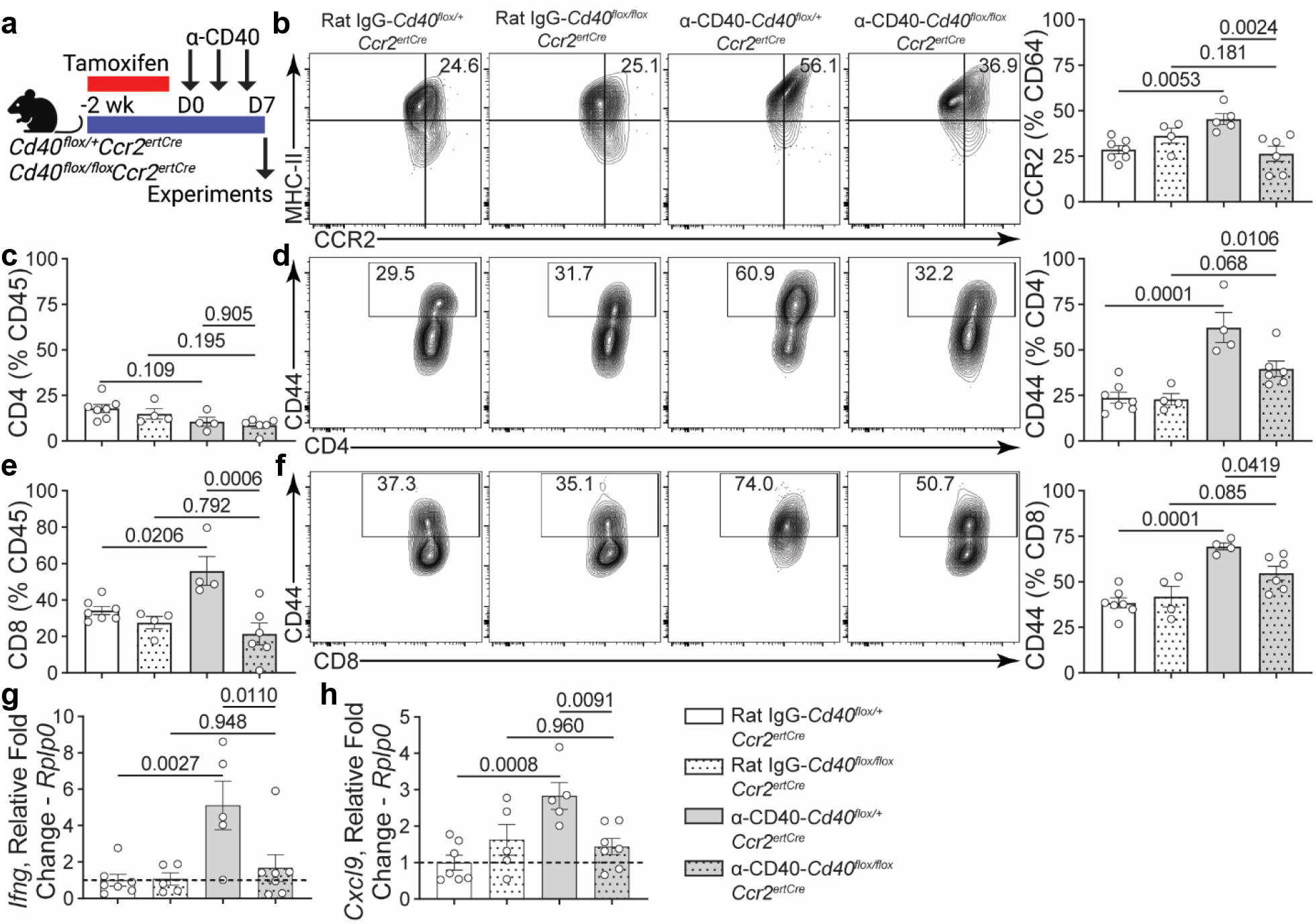
CD40 signaling in CCR2^+^ macrophages drives expansion and activation of CCR2^+^ macrophages and CD8 T-cells. **a,** Experimental model with mice receiving tamoxifen to conditionally delete CD40 expression in CCR2^+^ macrophages followed by α-CD40 agonist or Rat IgG isotype antibodies. **b,** Quantification of CCR2^+^ macrophages in the heart by flow cytometry in control (*Cd40^flox/+^Ccr2^ertCre^*) and CD40 deletion in CCR2 macrophages (*Cd40^flox/flox^Ccr2^ertCre^*) mice. Comparisons between Rat IgG-*Cd40^flox/+^Ccr2^ertCre^*(n = 7), Rat IgG-*Cd40^flox/flox^Ccr2^ertCre^* (n = 4), α-CD40-*Cd40^flox/+^Ccr2^ertCre^* (n = 5), and α-CD40-*Cd40^flox/flox^Ccr2^ertCre^* (n = 6). **c,** Quantification of CD4 T-cells, **d,** CD4^+^CD44^+^ T-cells, **e,** CD8 T-cells, and **f,** CD8^+^CD44^+^ T-cells cells in the heart by flow cytometry comparing Rat IgG-*Cd40^flox/+^Ccr2^ertCre^*(n = 7), Rat IgG-*Cd40^flox/flox^Ccr2^ertCre^* (n = 4), α-CD40-*Cd40^flox/+^Ccr2^ertCre^* (n = 4), and α-CD40-*Cd40^flox/flox^Ccr2^ertCre^* (n = 6). **g,** *Ifng*, and **h,** *Cxcl9* mRNA expression measured by RT-qPCR from bulk cardiac tissue in Rat IgG-*Cd40^flox/+^Ccr2^ertCre^*(n = 7), Rat IgG-*Cd40^flox/flox^Ccr2^ertCre^* (n = 5), α-CD40-*Cd40^flox/+^Ccr2^ertCre^* (n = 5), and α-CD40-*Cd40^flox/flox^Ccr2^ertCre^* (n = 7). One-way ANOVA with Sidak correction performed for all statistical analysis. Error bars indicate means ± s.e.m.

### Cardiac immune cell expansion occurs through an IFN-γ-dependent feedforward loop

Following anti-CD40 agonist antibody treatment, intracellular flow cytometry analysis demonstrated upregulation of IFN-γ in CD4 and CD8 T-cells with an effector memory phenotype (**Fig. 7a-b**). To determine the effect of neutralizing IFN-γ, wildtype mice received isotype control or IFN-γ antagonist antibody (i.p. 300 µg, single dose^25^) with concurrent anti-CD40 agonist antibody treatment (**Fig. 7c**). This resulted in a reduction of CCR2^+^ macrophages within the heart (**Fig. 7d**) and decreased *Ifng* and *Cxcl9* expression approaching baseline levels (**Fig. 7e-f**). We next crossed *Lyz2^Cre^* mice with *Ifngr1^flox/flox^*mice to generate mice lacking IFN-γ receptors in myeloid cells (including CCR2^+^ macrophages). Control and *Lyz2^Cre^Ifngr1^flox/flox^*mice were treated with isotype or anti-CD40 agonist antibodies (**Fig. 7g**). When compared to control littermates, *Lyz2^Cre^Ifngr1^flox/flox^*mice treated with anti-CD40 agonist antibodies had reduced CD68^+^ macrophages by immunostaining (**Fig. 7h**). Flow cytometry analysis confirmed a specific reduction of CCR2^+^ macrophages (**Fig. 7i**). Using *in situ* RNA hybridization for *Cxcl9*, we observed decreased expression of *Cxcl9* in *Lyz2^Cre^Ifngr1^flox/flox^* hearts compared to control hearts treated with anti-CD40 agonist antibodies (**Fig. 7j**). Lastly, *Ifng* expression was also significantly reduced in *Lyz2^Cre^Ifngr1^flox/flox^*hearts compared to control hearts treated with anti-CD40 agonist antibodies (**Fig. 7k**). These results demonstrate that anti-CD40 agonist antibodies increase IFN-γ producing T-cells and that IFN-γ signaling to macrophages is necessary for CCR2^+^ macrophage expansion and myocardial cytokine expression.

**Fig. 7.**
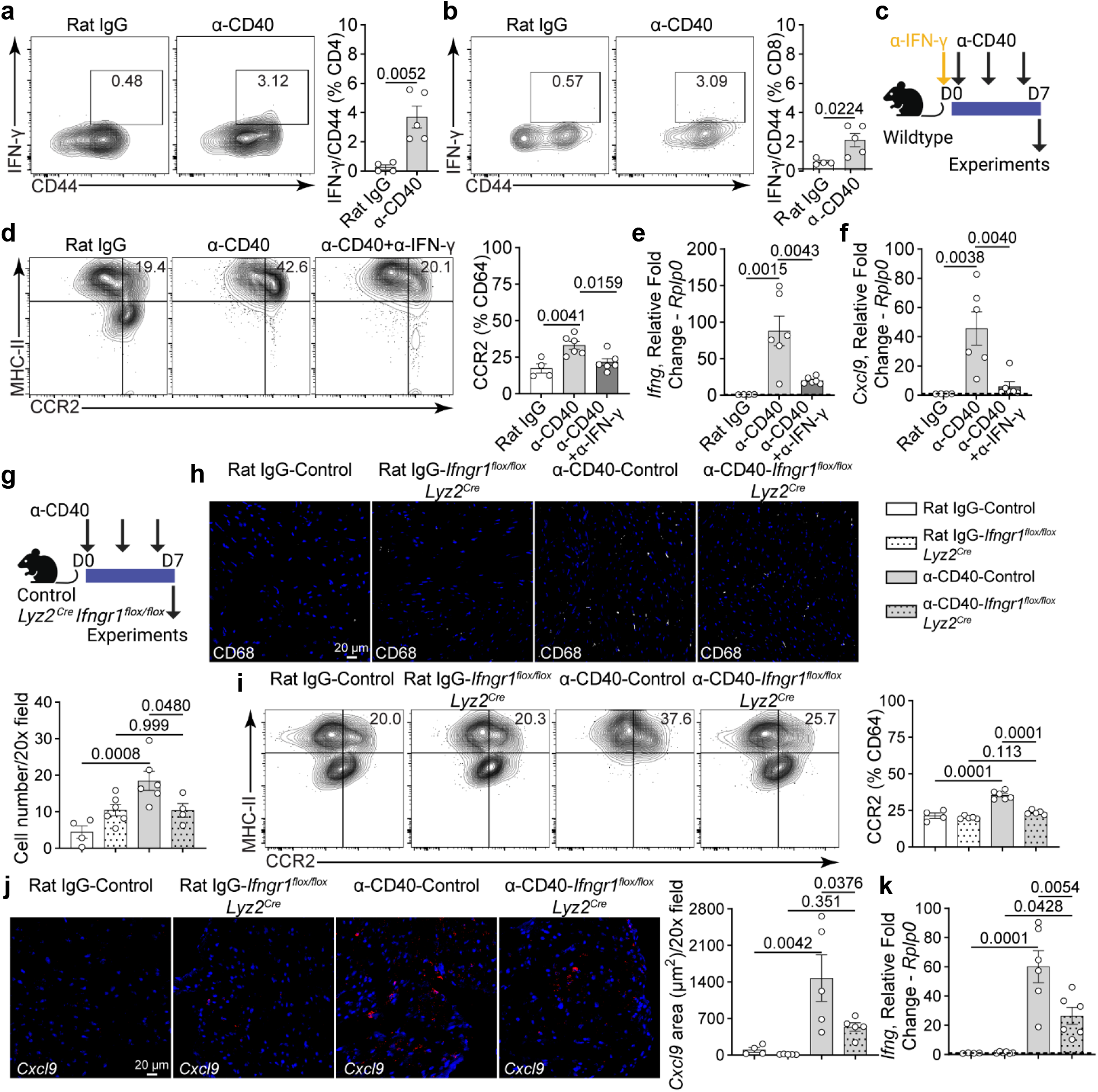
Cardiac immune cell expansion occurs through an IFN-γ-dependent feedforward loop. **a,** Quantification of IFN-γ^+^CD44^+^ CD4 T-cells and **b,** IFN-γ^+^CD44^+^ CD8 T-cells in the heart by flow cytometry between Rat IgG (n = 4) and α-CD40 (n = 5). Two-tailed unpaired Student’s *t*-test. **c,** Experimental model with mice receiving neutralizing INF-γ with concurrent α-CD40 agonist or IgG isotype antibodies. **d,** Quantification of CCR2^+^ macrophages in the heart by flow cytometry comparing Rat IgG (n = 4), α-CD40 (n = 6), and α-CD40+α-IFN-γ (n = 6). **e,** *Ifng*, and **f,** *Cxcl9* mRNA expression measured by RT-qPCR from cardiac tissue comparing Rat IgG (n = 4), α-CD40 (n = 6), and α-CD40+α-IFN-γ (n = 6). **g,** Experimental model with control or mice lacking IFN-γ receptors in myeloid cells (*Lyz2^Cre^Ifngr1^flox/flox^*) treated with α-CD40 agonist or Rat IgG isotype antibodies. **h,** Representative CD68 macrophage immunofluorescent staining (white) in Rat IgG-Control (n = 4), Rat IgG-*Lyz2^Cre^Ifngr1^flox/flox^* (n = 6), α-CD40-Control (n = 6), and α-CD40-*Lyz2^Cre^Ifngr1^flox/flox^*(n = 4) hearts, and quantification of CD68^+^ cells per 20x field. Scale bar 20 µm. **i,** Quantification of CCR2^+^ macrophages in the heart by flow cytometry comparing Rat IgG-Control (n = 4), Rat IgG-*Lyz2^Cre^Ifngr1^flox/flox^*(n = 5), α-CD40-Control (n = 6), and α-CD40-*Lyz2^Cre^Ifngr1^flox/flox^*(n = 5). **j,** Representative images of *Cxcl9* (red) expression detected in Rat IgG-Control (n = 4), Rat IgG-*Lyz2^Cre^Ifngr1^flox/flox^* (n = 5), α-CD40-Control (n = 5), and α-CD40-*Lyz2^Cre^Ifngr1^flox/flox^* (n = 5) mouse hearts by RNA *in situ* hybridization. Scale bar 20 µm. Quantification of *Cxcl9* fluorescent area per 20x field. **k,** *Ifng* mRNA expression measured by RT-qPCR from bulk cardiac tissue comparing Rat IgG-Control (n = 4), Rat IgG-*Lyz2^Cre^Ifngr1^flox/flox^* (n = 6), α-CD40-Control (n = 6), and α-CD40-*Lyz2^Cre^Ifngr1^flox/flox^* (n = 6). One-way ANOVA with Sidak correction performed for all statistical analysis except as noted. Error bars indicate means ± s.e.m.

We next focused on the mechanism by which CD40 signaling in CCR2^+^ macrophages drives T-cell IFN-γ production. We tested the possibility that CD40 mediated induction of IL-12b and TNF in CCR2^+^ macrophages signals to T-cells to trigger IFN-γ production. Wildtype mice were exposed to anti-CD40 agonist antibodies and were treated with either isotype control or concurrent anti-IL12b and anti-TNF neutralizing antibodies (250 µg every 72 hours)^26, 27^ (**Fig. 8a**). Immunostaining revealed reduction of CD8 T-cells in the heart following neutralization of IL12b and TNF, compared to isotype control (**Fig. 8b**). In addition, there was decreased IFN-γ production in CD8 T-cells with an effector memory phenotype in mice receiving concurrent neutralization of IL12b and TNF antibodies (**Fig. 8c**). Neutralization of IL12b or TNF alone did not have an effect on IFN-γ production from effector memory CD8 T-cells (**Extended Data Fig. 7a-b**).

**Fig. 8.**
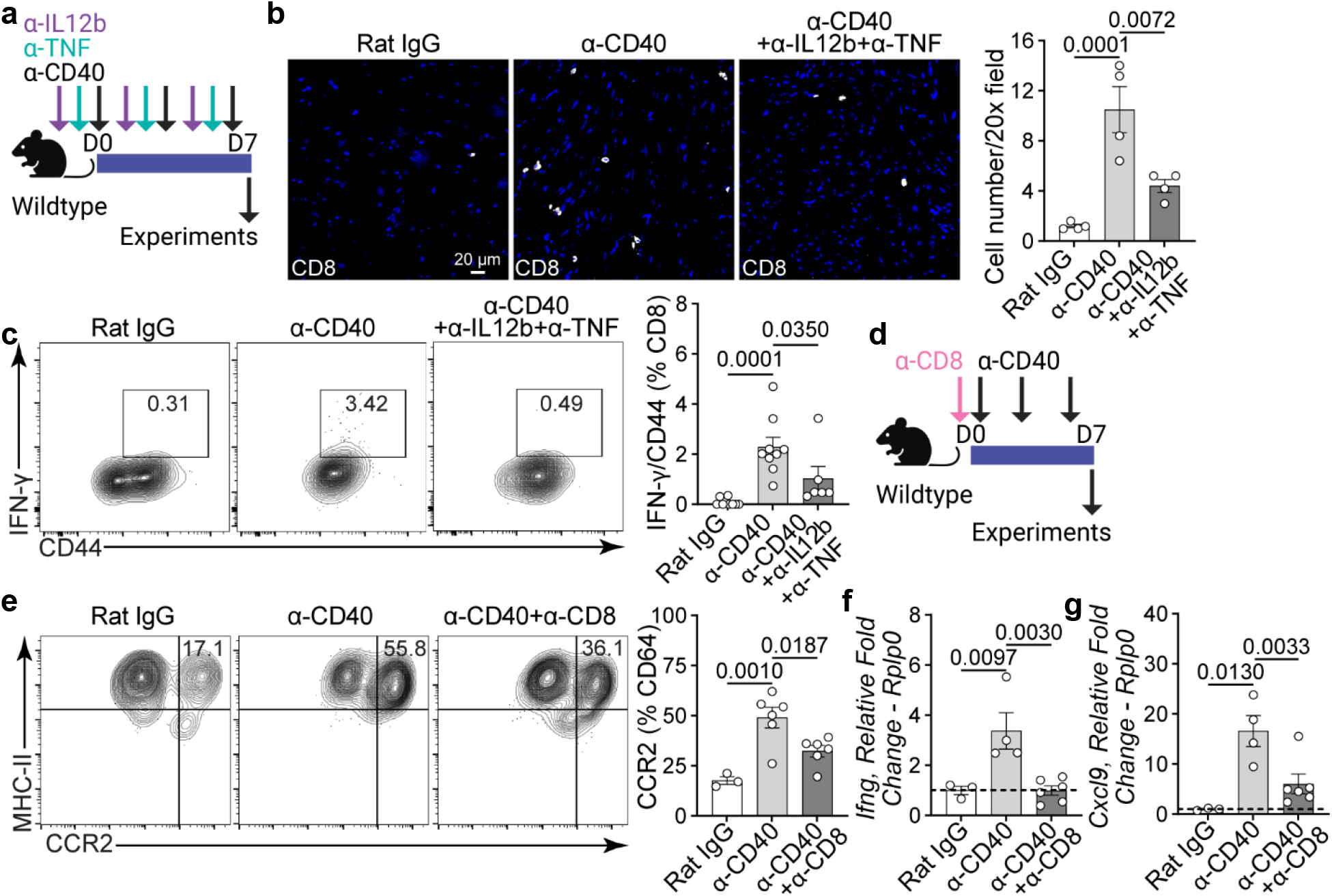
IL12b and TNF regulate CD8 T-cell IFN-γ production. **a,** Experimental model with mice receiving neutralizing α-IL12b and α-TNF with concurrent α-CD40 agonist or Rat IgG isotype antibodies. **b,** Representative CD8 T-cell immunofluorescent staining (white) in Rat IgG (n = 4), α-CD40 (n = 4), and α-CD40+α-IL12b+α-TNF (n = 4) hearts, and quantification of CD8^+^ T-cells per 20x field. Scale bar 20 µm. **c,** Quantification of IFN-γ^+^CD44^+^ CD8 T-cells in the heart by flow cytometry between Rat IgG (n = 9), α-CD40 (n = 9), and α-CD40+α-IL12b+α-TNF (n = 5). Data collected from 2 independent experiments. **d,** Experimental model with mice receiving depleting α-CD8 with concurrent α-CD40 agonist or Rat IgG isotype antibodies. **e,** Quantification of CCR2^+^ macrophages in the heart by flow cytometry comparing Rat IgG (n = 3), α-CD40 (n = 6), and α-CD40+α-CD8 (n = 6). **f,** *Ifng*, and **g,** *Cxcl9* mRNA expression measured by RT-qPCR from cardiac tissue in Rat IgG (n = 3), α-CD40 (n = 4), and α-CD40+α-CD8 (n = 6). One-way ANOVA with Sidak correction performed for all statistical analysis. Error bars indicate means ± s.e.m.

To establish a causal relationship between CD8 T-cells and CCR2^+^ macrophage activation, wildtype mice were stimulated with anti-CD40 agonist antibody and concurrently treated with either isotype or CD8 depleting antibody (i.p. 100 µg, single dose)^28^ (**Fig. 8d**). We observed decreased CCR2^+^ expansion in the heart following CD8 T-cell depletion (**Fig. 8e**) with concomitant decreased expression of *Ifng* and *Cxcl9* (**Fig. 8f-g**). CD8 T-cell depletion was confirmed by flow cytometry of splenocytes (**Extended Data Fig. 8a**). Depletion of CD4 T-cells alone had minimal impact on CCR2^+^ expansion in anti-CD40 agonist antibody treated mice. Furthermore, CD4 T-cell depletion had no additive effect when combined with CD8 T-cell depletion (**Extended Data Fig. 8b**). These findings delineate a pathway by which CD40 signaling in CCR2^+^ macrophages promotes the expression of TNF and IL12b, which signal to CD8 T-cells to produce IFN-γ resulting in the ongoing activation and expansion of CCR2^+^ macrophages.

### CD40 agonist treatment accelerates LV remodeling

To determine the effect anti-CD40 agonist antibody treatment has on LV remodeling, two clinically relevant models of cardiac injury were used: hypertensive cardiac fibrosis and reperfused myocardial infarction models. For both models, WT mice were treated with either isotype or anti-CD40 agonist antibodies every 72 hours for a total 35 days. In the first injury model, myocardial injury was induced by continuous infusion of angiotensin II and phenylephrine (AngII/PE)^29^ (**Fig. 9a**). We observed increased fibrosis by Masson’s trichrome staining in heart sections (**Fig. 9b**). Immunohistochemistry revealed increased CD45^+^ immune cell abundance (**Fig. 9c**). CD40 agonism resulted in upregulated gene expression of multiple inflammatory cytokines including *Ifng*, *l12b*, and *Tnf* and chemokines including *Ccl5*, *Cxcl9*, and *Cxcl10* at day 7 (**Fig. 9d**). To determine whether a similar response was evident in a second cardiac injury model, isotype control or anti-CD40 agonist antibody treated mice underwent reperfused myocardial infarction (**Fig. 9e**)^30^. 4 weeks after MI, we observed increased infarct size, reduced ejection fraction, and increased (worse) wall motion score index in mice treated with anti-CD40 agonist antibody (**Fig. 9f-j**). Collectively these data demonstrate that CD40 agonism sensitizes the heart to secondary cardiac injury resulting in increased cardiac inflammation, fibrosis, and LV remodeling.

**Fig. 9.**
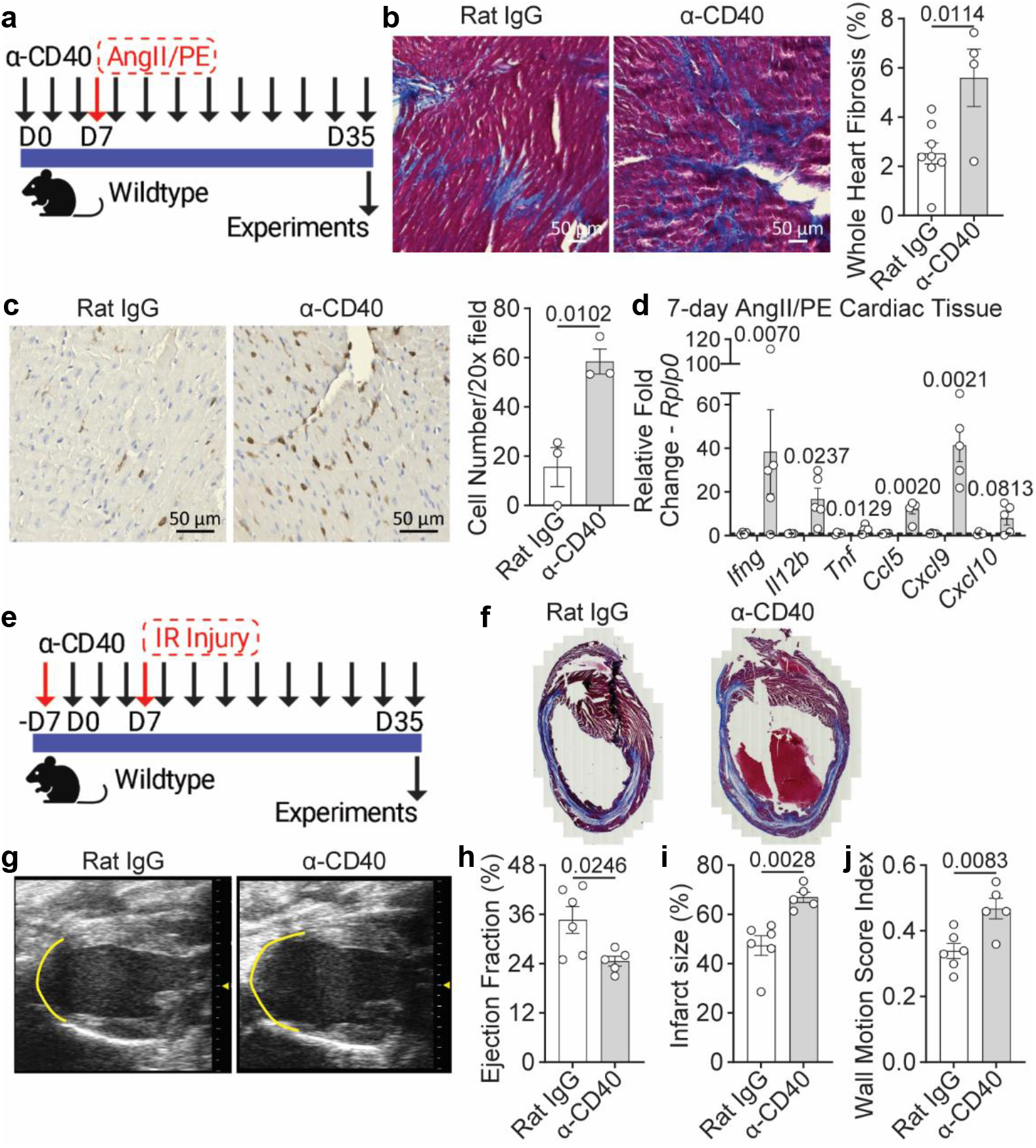
CD40 agonist treatment accentuates cardiac injury and remodeling. **a,** Experimental model with myocardial injury induced in wildtype mice by continuous infusion of angiotensin II and phenylephrine (AngII/PE) and concurrent treatment with α-CD40 agonist or Rat IgG isotype antibodies every 72 hours for a total of 35 days. **b,** Representative Masson’s trichrome staining showing myocardial fibrosis (blue staining) in Rat IgG (n = 8) and α-CD40 (n = 4) hearts and quantification of whole heart fibrosis. Scale bar 50 µm. **c,** Representative CD45 cell immunohistochemical staining (brown) in Rat IgG (n = 3) and α-CD40 (n = 3) in hearts, and quantification of CD45^+^ cells per 20x field. Scale bar 50 µm. **d,** *Ifng*, *Il12b*, *Tnf*, *Ccl5*, *Cxcl9*, and *Cxcl10* mRNA expression measured by RT-qPCR from 7-day AngII/PE bulk cardiac tissue. Comparison between Rat IgG (n = 4) and α-CD40 (n = 5) for each individual gene. **e,** Experimental model with myocardial injury in wildtype mice that underwent reperfused myocardial infarction injury and concurrent treatment with α-CD40 agonist or Rat IgG isotype antibodies every 72 hours for a total of 35 days. **f,** Representative Masson’s trichrome staining showing short axis cross sections with myocardial fibrosis (blue staining) in Rat IgG and α-CD40 hearts. **g,** Representative 2-dimensional echocardiography showing parasternal long axis images with outline of infarct (yellow line) in Rat IgG and α-CD40 hearts. **h,** Quantification of % ejection fraction, **i,** % infarct size, and **j,** wall motion score index in Rat IgG (n = 6) and α-CD40 (n = 5) hearts. Two-tailed unpaired Student’s *t*-test performed for all statistical analysis. Error bars indicate means ± s.e.m.

## Discussion

Adverse cardiac events following cancer immunotherapy vary from transient arrhythmias to fulminant and life-threatening myocarditis^31, 32^. While inhibition of conventional ICI signaling (PD-1, PD-L1, CTLA-4) is associated with infiltration of macrophages, CD8 and CD4 T-cells into the heart, much less is known regarding the potential for novel immune checkpoint modulators to trigger cardiac inflammation. This represents a critical question for the field as anti-CD40 agonist antibodies and anti-LAG-3 neutralizing antibodies are increasingly being used alone or in combination with conventional ICIs to treat previously uncurable cancers.

Here, we used various genetic mouse models in conjunction with immune profiling, single cell RNA sequencing, antibody neutralization/depletion studies, and cardiac injury models to demonstrate that anti-CD40 agonist antibodies trigger cardiac inflammation and sensitize the heart to subsequent cardiac injury. Mechanistically, anti-CD40 agonist antibodies activate signaling in cardiac CCR2^+^ macrophages leading to production of cytokines (IL12b, TNF) that activate CD8 T-cells to produce IFN-γ. CCR2^+^ macrophages represent a key target of IFN-γ signaling, which drives CCR2^+^ macrophage expansion and initiation of a positive feedback loop between CD8 T-cells and CCR2^+^ macrophages potentially through CXCL9 and CCL5 signaling. Cardiac inflammation could be inhibited by deleting CD40 or IFNGR1 in CCR2^+^ macrophages, depleting CD8 T-cells, neutralization of IFN-γ signaling or TNF and IL12b signaling. From a clinical standpoint, exposure to anti-CD40 agonist antibody treatment potentiated LV remodeling in response to AngII/PE infusion or reperfused MI.

Unlike anti-PD-1, anti-PD-L1, and anti-CTLA-4 antibodies that function by inhibiting T-cell activation, anti-CD40 agonist antibodies directly activate antigen presenting cells resulting in cytokine production, T-cell priming, activation, and proliferation. Macrophages represent the major antigen presenting cell in the heart. Among cardiac macrophage subsets, CCR2^+^ macrophages appear to be the primary target of anti-CD40 agonist antibodies. CD40 agonism triggers expression of pro-inflammatory chemokines and cytokines (CXCL9, CCL5, IL12b, TNF) from CCR2^+^ macrophages and expansion of this cell population through a combination of local proliferation and monocyte recruitment. In studies investigating ICI associated adverse events including human ICI myocarditis or colitis, macrophages expressing CXCL9 have also been identified^33, 34^. CXCL9 is known to be induced by type II interferon signaling and contributes to a CXCL9/10-CXCR3 axis that facilitates interactions between antigen presenting cells and T-cells resulting in the generation of effector CD8 T-cells^35^. The initiation and maintenance of this inflammatory amplification loop of CCR2^+^ macrophages and CD8 T-cells mediated through IFN-γ is of considerable interest as it may provide an opportunity to disrupt cardiac immune cell expansion and inflammation that is seen in ICI myocarditis.

Adverse cardiac events remain a major challenge for cancer immunotherapies. We identified several potential therapeutic approaches that may disrupt the inflammatory loop of activation generated by CD40 and IFN-γ signaling in the heart. Collectively, these strategies converge on interrupting IFN-γ signaling, but it remains to be determined what effect this approach will have on the therapeutic efficacy for tumor control. While conflicting reports are present in the literature, some studies have suggested that IFN-γ signaling may contribute to tumor control by inducing apoptosis, necroptosis, and senescence^36–38^. Thus, it is imperative to understand whether inhibition of IFN-γ signaling in the tumor microenvironment impacts tumor control in the setting of immune checkpoint therapies. One possible approach to avoid immune related adverse events of immune checkpoint therapies is to minimize systemic CD40 and IFN-γ activation. This may be achieved by direct intratumoral injection of immune checkpoint therapeutics or use of bispecific agonist anti-CD40 antibodies targeted to the tumor microenvironment through tumor specific antigens^39, 40^. Such strategies to minimize systemic toxicity particularly in the heart are of considerable interest, since adverse cardiac events such as ICI myocarditis carry the highest mortality risk and which upon diagnosis, results in discontinuation of life saving cancer immunotherapies.

We acknowledge that this study is not without limitations. CD40 signaling is a well-established activator of dendritic cells. Our data showed a modest expansion of classical dendritic cells in the heart using *Zbtb* GFP-reporter mice. As such we cannot exclude a role for classical dendritic cells. Interestingly, prior studies focused on autoimmune myocarditis similarly demonstrated limited expansion of dendritic cells in the heart. However, inhibition of dendritic cells resulted in only a modest reduction in the myocarditis phenotype^41^. Formal dendritic cell depletion studies are required to address the exact role for dendritic cells in cardiac inflammation elicited by CD40 signaling activation. This is of importance as considerable attention has been focused on generating bispecific antibodies that preferentially activate dendritic cells following CD40 activation^42^, which may serve as a valuable strategy to prevent broader CD40 activation in CCR2^+^ macrophages that may lead to adverse cardiovascular events. Of note, we observed no evidence of B cell expansion or involvement in CD40 triggered myocardial inflammation. Future work will focus on the impact of concomitant CD40 agonism and PD-1 or PD-L1 blockade on cardiac inflammation as well as immune training elicited by different tumor types. Investigating tumor bearing models will be essential to ascertain the effect of strategies meant to suppress cardiac inflammation on short-term and long-term tumor control.

In conclusion, this study demonstrates that anti-CD40 agonist antibody exposure elicits cardiac inflammation by triggering a feed-forward inflammatory loop between cardiac macrophages and CD8 T-cells and sensitizes the heart to subsequent injury resulting in accelerated LV remodeling. These findings identify a possible cardiovascular risk of CD40 agonists.

## Acknowledgements

JJ is supported by funding provided by the National Institutes of Health (1K08HL163518-01A1, R25 HL105400), the Paul and Patti Eisenberg Scholar Award of Washington University in the Division of Cardiology, and Institutional Research Grant IRG-21-133-64 from the American Cancer Society. KJL is supported by the Washington University in St. Louis Rheumatic Diseases Research Resource-Based Center grant (NIH P30AR073752), the National Institutes of Health [R01 HL138466, R01 HL139714, R01 HL151078, R01 HL161185, R35 HL161185], Leducq Foundation Network (#20CVD02), Burroughs Welcome Fund (1014782), and Children’s Discovery Institute of Washington University and St. Louis Children’s Hospital (CH-II-2015-462, CH-II-2017-628, PM-LI-2019-829), Foundation of Barnes-Jewish Hospital (8038-88), and generous gifts from Washington University School of Medicine. We are grateful to Dr. Esther Lutgens (Amsterdam University Medical Centre) for providing us with *Cd40^flox/flox^* mice. We are thankful to the Mouse Cardiovascular Phenotyping Core facility at Washington University for performing mouse echocardiography and the reperfused myocardial infarction surgeries. We are thankful to the Genome Technology Access Center at the McDonnell Genome Institute for help with genomic analysis, the Digestive Diseases Research Core Center (DDRCC) for histology services and the Washington University Center for Cellular Imaging (WUCCI) at Washington University School of Medicine.

## Author Contributions

JJ, JA, and KJL were responsible for conceptualization of the study. JJ, JA, PM, XW, RD, and KJL contributed to experimental design, acquired data, and performed data analysis. JJ and JA performed statistical analysis. JJ and KJL wrote the original draft of the manuscript, and all authors approved the final version.

## Competing Interests

The authors declare no competing interests.

## Methods

### Animal Models

All experimental animal procedures were performed in accordance with IUCAC approved protocols. Mice were purchased or bred, then maintained at the Washington University School of Medicine. Mouse strains utilized include: C57BL/6J mice (The Jackson Laboratory, 000664), *Ccr2^gfp/+^* ^43^, *Zbtb46^gfp/+^* ^14^, *Ccr2^ertCre^* ^44^, *Rosa26^tdTomato^* ^45^*, Cd40^flox/flox^* ^23^, *Lyz2^Cre^* ^46^, *Ifngr1^flox/flox^* ^47^. All genetic mice were on the C57BL/6J background and genotyped according to established protocols. Experiments were performed on mice 2-8 months age. Similar numbers of male and female mice were used for experiments. For Cre recombination, mice received i.p. injections of 60 mg/kg tamoxifen (Millipore Sigma, T5648) every 3 days for 1 week in *Ccr2^ertCre^Rosa26^tdTomato^* and i.p. injections of 40 mg/kg tamoxifen daily for 5 days followed by tamoxifen food pellets (Envigo, TD.130857) for 2 weeks in *Cd40^flox/flox^Ccr2^ertCre^* mice.

### Antibody Treatment

Mice received i.p injections with the following antibodies: isotype control 100 µg Rat IgG (clone 2A3); agonist 100 µg α-CD40 (clone FGK4.5/FGK45); neutralizing 300 µg α-IFN-γ (clone R4-6A2); neutralizing 250 µg α-TNF (clone TN3-19.12); neutralizing 250 µg α-IL12b (clone R2-9A5); B cell depleting 200 µg α-CD19 (clone 1D3); CD8 depleting 100 µg α-CD8 (clone 2.43); and CD4 depleting 200 µg α-CD4 (clone GK1.5). All antibodies were purchased from Bio X Cell (bioxcell.com). In all experiments, mice received either Rat IgG or α-CD40 for 7, 28, or 35 days every 72 hrs. In the IFN-γ neutralizing studies, mice received a single dose of α-IFN-γ at the start of the experiment. In the IL12b and TNF neutralizing studies, mice received α-IL12b and/or α-TNF every 72hrs at the start of the experiment. In the B cell depletion studies, mice received α-CD19 three days prior to receiving the first dose of concurrent Rat IgG or α-CD40, then received a second dose 72 hrs later for a total of 2 doses. In the CD8 and CD4 T-cell depletion studies, mice received single doses of α-CD8 and/or α-CD4 at the start of the experiment.

### Histology

At experiment termination, mouse hearts were perfused with cold PBS, fixed overnight with 4% PFA in PBS at 4⁰C, then placed into histology cassettes for paraffin embedding and dehydration in 70% ethanol in water. Hearts were paraffin embedded, cut into 5 µm sections, and mounted on positively charged slides. H&E or Masson’s trichrome tissue staining was completed using standard published protocols. Whole tissue digital images were taken using a Zeiss AxioScan Z1 slide scanner with10x objective and analyzed using Zen 2.6 (Blue Edition). For fibrosis analysis, the area of fibrosis was detected in the whole heart section for each condition using ImageJ software (version 1.53r). For CD45 staining, paraffin-embedded sections were dewaxed in xylene, rehydrated, endogenous peroxide activity quenched in 10% methanol and 3% hydrogen peroxide, processed for antigen retrieval by boiling in citrate buffer pH 9.0 for 15 minutes, then blocked in 10% bovine serum albumin containing 0.05% Tween-20, and stained with 1:100 primary antibody CD45 (BD Bioscience, 550539) overnight at 4 °C: The primary antibody was detected using biotinylated anti-rat secondary antibody (Vector Laboratories, BA40001.5) in conjunction with streptavidin horseradish peroxidase (ABC Elite, Vector Labs). The DAB Substrate Peroxidase Kit (SK-4100, Vector Labs) was utilized to visualize antibody staining per manufacturer protocol. At least 10 views were randomly selected to analyze the number of positively stained cells in the heart and displayed as an average cell number per 20x field for each condition.

### Immunofluorescence

At experiment termination, mouse hearts were perfused with cold PBS and fixed overnight at 4⁰C with 4% PFA in PBS, followed by dehydration in 30% sucrose in PBS overnight at 4⁰C, then embedded in O.C.T. compound and frozen in -80°C per standard protocol. To generate fixed frozen slides, hearts were sectioned at 10 µm using a Leica Cryostat. Slides were brought to room temperature for 5 minutes and washed in Tris-Buffered Saline (TBS) for 5 minutes. Sections were permeabilized in 0.05% Tween20 in TBS (TBST) for 5 minutes followed by blocking with 10% Fetal Bovine Serum (FBS) in TBST for 30 min at room temperature. Sections were stained with primary antibody diluted in 10% FBS in TBST overnight at 4°C. Primary antibodies used: CD68 (clone FA-11, BioLegend 137002) 1:200 dilution; CD8 (clone EPR21769, Abcam, ab217344) 1:200 dilution; GFP (Abcam, ab13970) 1:1000 dilution; Ki67 (clone SP6, Abcam, ab16667) 1:250 dilution. Sections were washed in TBS then incubated with appropriate immunofluorescent secondary antibodies at 1:500 dilution in 10% FBS in TBST for 60 min at room temperature. Slides were washed with TBST, mounted with DAPI mounting media (Millipore Sigma, F6057), and cover slipped. Slides were stored at 4⁰C, protected from light, and imaged within 3 days. Immunofluorescence staining of paraffin embedded tissues were processed using the OPAL 4-Color Manual IHC Kit (Akoya Biosciences, NEL810001KT).

Primary antibodies used: CD64 (R&D Systems, AF2074) 1:200 dilution; CD8 (clone EPR21769, Abcam, ab217344) 1:200 dilution. Immunofluorescence staining of fixed frozen slides were also processed for *in situ* RNA hybridization using the RNAscope Multiplex Fluorescent Detection Kit v2 (ACD, 323110) with probes for *Cxcl9* (ACD, 489341) and *Ccr2* (ACD, 433271). All slides were imaged using a Zeiss LSM 700 confocal microscope with a 20x objective and analyzed using Zen 2.6 (Blue Edition). At least 4-5 distinct sections were quantified to analyze the number of positively stained cells in the heart and displayed as an average cell number per 10x or 20x field for each condition. For quantification of *in situ* hybridization, 4-5 distinct sections were quantified to analyze percentage of fluorescence in the heart for each condition.

### Flow Cytometry

At experiment termination, single cell suspensions of mouse hearts were generated following perfusion with cold PBS, atria removal, fine mincing, and digestion for 20 minutes at 37⁰C in DMEM (Gibco, 11965-084) containing 4500 U/ml Collagenase IV (Millipore Sigma, C5138), 2400 U/ml Hyaluronidase I (Millipore Sigma, H3506), and 6000 U/ml DNAse I (Millipore Sigma, D4527). Enzymes were deactivated with HBSS (Gibco, 14025092) containing 2% FBS and 0.2% BSA, then filtered through 40-micron strainers. For red blood cell lysis, cells were incubated with ACK lysis buffer (Gibco, A10492-01) for 5 minutes at room temperature. Cells were washed with DMEM and resuspended in FACS buffer (PBS containing 2% FBS and 2 mM EDTA). Cells were incubated in a 1:200 dilution of fluorescence conjugated monoclonal antibodies (see list of antibodies below) for 30 minutes at 4⁰C in the dark. Samples were washed in FACS buffer and resuspended in a final volume of 300 µl FACS buffer. For splenocyte single cell suspensions, spleens were homogenized by using a flat plunger end to push spleen through a 40-micron strainer and into a container with DMEM. Cells were then processed with ACK lysis buffer and processed with remaining steps as already described. For intracellular staining of IFN-γ, mice received an i.p. injection with 150 µg Brefeldin A Solution (BioLegend, 420601) in PBS before hearts were processed for single cell suspension^48^. Following staining with surface markers, cells were processed with BioLegend Cyto-Fast Fix/Perm Buffer Set (BioLegend 426803), stained with IFN-γ, and resuspended in 250 µL FACS buffer. Flow cytometry and sorting was performed using a BD FACSMelody Cell Sorter (BD Biosciences). Flow analysis was performed using FlowJo (v10.10.0 for Windows). Depending on immune cell panels (myeloid or lymphoid) and transgenic mice with fluorescent markers (GFP or tdTomato), FACS was performed using the following antibodies:

CCR2 BV421 (BioLegend, 150605)

CD19 PE (BioLegend, 115508)

CD4 PE/Cy7 (BioLegend, 100422) or FITC (BioLegend, 100406) or BV510 (BioLegend, 100559)

CD44 APC (BioLegend, 103012)

CD45 PerCP/Cy5.5 (BioLegend, 103132) or BV510 (BioLegend, 103138)

CD64 APC (BioLegend, 139306) or PE-Cy7 (BioLegend, 139313)

CD8 APC/Cy7 (BioLegend, 100721) or PerCP/Cy5.5 (BioLegend, 100734)

DAPI (BD Pharmingen, 564907)

IFN-γ PE (BD Biosciences, 554412)

Ly6g FITC (BioLegend, 127606) or APC/Cy7 (BioLegend, 127623)

Ly6c PE/Cy7 (BioLegend, 128017) or BV510 (BioLegend, 128033)

1A/1E (MHCII) APC/Cy7 (BioLegend, 107628)

### Real Time Quantitative Polymerase Chain Reaction

Mouse heart tissue that was flash frozen in liquid nitrogen then stored in -80°C was processed using TRIzol-chloroform (Invitrogen, 15596026), stainless steel beads (Qiagen, 69989), and TissueLyser LT (Qiagen) per manufacturer methods. RNA was isolated using the PureLink PCR Purification Kit (Invitrogen, K310001). RNA concentration was measured using a nanodrop spectrophotometer (ThermoFisher Scientific). cDNA was generated using the High-Capacity cDNA Reverse Transcription Kit (Applied Biosystems, 4368814). RT-qPCR reactions were prepared with sequence-specific primers (Integrated DNA Technologies, idtdna.com) with Power SYBR Green PCR Master Mix (Applied Biosystems, 4367659). RT-qPCR was performed using a QuantStudio 6 Flex Real-Time PCR System (Thermo Fisher Scientific) per manufacturer methods and as previously published^49^. *Rplp0* was used as reference control and all individual gene expression was normalized to *Rplp0* expression.

*Ccl5* Forward: 5’-GGG TAC CAT GAA GAT CTC TGC-3’ Reverse: 5’-TCT AGG GAG AGG TAG GCA AAG-3’

*Cxcl9* Forward: 5’-GTT CGA ACC CTA GTG ATA AG-3’ Reverse: 5’-GTT TGA GGT CTT TGA GGG ATT TG-3’

*Cxcl10* Forward: 5’-TCA GGC TCG TCA GTT CTA AGT-3’ Reverse: 5’-CCT TGG GAA GAT GGT GGT TAA G-3’

*Ifng* Forward: 5’-GGC CAT CAG CAA CAA CAT AAG-3’ Reverse: 5’-GTT GAC CTC AAA CTT GGC AAT AC-3’

*Il12b* Forward: 5’-GCA GAT GAA GCC TTT GAA GAA C-3’ Reverse: 5’-AGA ACT TGA GGG AGA AGT AGG A-3’

*Rplp0* Forward: 5’-ATC CCT GAC GCA CCG CCG TGA-3’ Reverse: 5’-TGC ATC TGC TTG GAG CCC ACG TT-3’

*Tnf* Forward: 5’-CTT CTG TCT ACT GAA CTT CGG G-3’ Reverse: 5’-CAG GCT TGT CAC TCG AAT TTT G-3’

### CITE-Seq Analysis

Single cell suspensions were prepared and pooled from 4 hearts for fluorescence activated cell sorting from C57BL/6J mice following α-CD40 agonist or Rat IgG treatment for 7 days. CD45^+^ cells (10,000-15,000 cells) were collected for CITE-Seq and processed using the 10X Single Cell Gene Expression v3.1 kit (prepares cDNA libraries for gene expression and cell surface protein analysis) that were sequenced at a depth of 75,000 reads/cell (Nova-seq) at the McDonnel Genome Institute (MGI). Downstream data processing was performed in the R package Seurat (v4) 57. SCtransform was used to scale and normalize the data for RNA counts. Quality control filters of genes/cell, mitochondrial reads/cell, and read counts/cell were applied. PCA was used for dimensionality reduction, data was clustered at multiple resolutions, and UMAP projections were generated for data visualization. Differential gene expression of upregulated genes in each subpopulation was generated in Seurat using FindAllMarkers with a minimum fraction (min.pct) of 0.1 and a logFC threshold of 0.1. Weighted nearest neighbor analysis using Seurat v4 was performed to integrate protein and RNA information in T/NK cell compartment. For pathway analysis and volcano plots, genes with an adjusted p-value < 0.05 and absolute(log2FC > 1.0) were deemed significant and analyzed using Enrichr software (maayanlab.cloud/Enrichr).

### Hypertensive Cardiac Fibrosis Injury

For hypertensive cardiac fibrosis injury, 2-month-old C57BL/6J mice underwent subcutaneous implantation of Alzet Osmotic Pumps (Alzet) on their back as per manufacturer guidelines. Minipumps were filled and implanted to continuously secrete 1.5 μg/g/day angiotensin II (Bachem, 4006473) and 50 μg/g/day phenylephrine (Millipore Sigma, P6126-5G) or 0.9% normal saline for 7 (Alzet, 2001) or 28 (Alzet, 2004) days. Hearts were harvested to generate paraffin embedded slides as already described and processed for immunohistopathology. For 7-day AngII/PE studies, hearts were processed for gene expression analysis as already described.

### Reperfused Myocardial Infarction

Surgery was performed in the Washington University Mouse Cardiovascular Phenotyping Core facility as previously described^7, 50^. Briefly, 2-month-old C57BL/6J mice were anesthetized, intubated, and mechanically ventilated. The heart was exposed, and a suture placed around the proximal left coronary artery. The suture was threaded through a 1 mm piece of polyethylene tubing to serve as the arterial occlude. Each end of the suture was exteriorized through the thorax. The skin was closed, and mice were given a 2-week recovery period prior to induction of ischemia. After 2 weeks, the animals were anesthetized and placed on an Indus Mouse Surgical Monitor system to accurately record ECG during ischemia. Ischemia was induced after anesthetizing the animals. Tension was exerted on suture ends until ST-segment elevation was seen via ECG. Following 90 minutes of ischemia time, tension was released, and the skin was then closed. Hearts were harvested to generate paraffin embedded slides as already described and processed for immunohistopathology.

### Echocardiography

At the end of the reperfused myocardial infarction studies, mice underwent echocardiography, which was performed in the Washington University Mouse Cardiovascular Phenotyping Core facility as previously described^7, 50^. Briefly, mice were sedated with Avertin (0.005 ml/g) and 2D and M-mode images were obtained in the long and short axis views 4 weeks after reperfused myocardial infarction using the VisualSonics770 Echocardiography System. Quantification of left ventricular internal diameter during diastole, left ventricular internal diameter during systole, % ejection fraction, infarct size, and wall motion score index were measured using edge detection and tracking through Vevo LAB software (v3.1.1). Percent infarct size was determined by measuring the perimeter of the akinetic portion of the LV myocardium and normalizing it to the perimeter of the total LV myocardium.

### Statistical Analysis

Analysis of 2 groups for histology, immunohistochemistry, immunofluorescence, flow cytometry, RT-qPCR, *in situ* RNA hybridization, and echocardiography experiments were analyzed using two tailed unpaired Students *t*-test assuming equal variance. Analysis of 3 groups or more were analyzed using one-way ANOVA with Sidak correction. P < 0.05 was a priori considered statistically significant. Analysis and plotted data were completed using GraphPad Prism (version 10.1.2 for Windows).

## Data Availability

CITE-sequencing data will be available on the Gene Expression Omnibus at the time of publication.

## Code Availability

R scripts are available on reasonable request.

## Notes

### Competing Interest Statement

The authors have declared no competing interest.

