## Extended Data Figures for "CD40 is an immune checkpoint regulator that potentiates myocardial inflammation through activation and expansion of CCR2^+^ macrophages and CD8 T-cells"

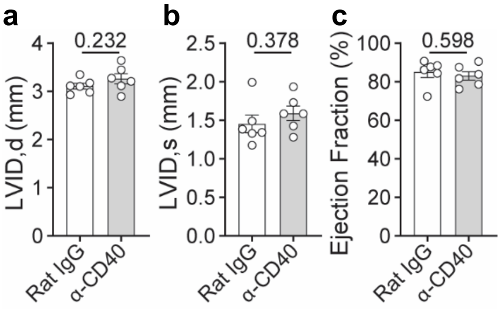


**Extended Data Fig. 1. No change in baseline heart chamber size or function in CD40 agonist treated mice. a,** Quantification of left ventricular internal diameter during diastole (LVID,d), **b,** left ventricular internal diameter during systole (LVID,s), and **c,** % ejection fraction using 2-dimensional echocardiography in mice treated with Rat IgG (n = 6) or α-CD40 (n = 6) for 28 days. Two-tailed unpaired Student's *t*-test performed for all statistical analysis. Error bars indicate means ± s.e.m.

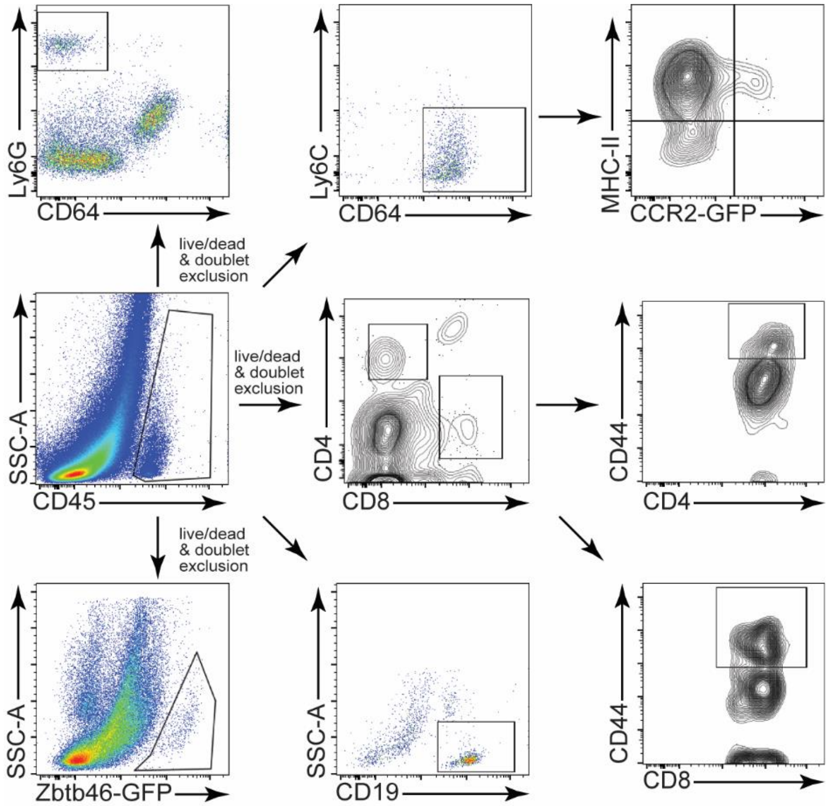


**Extended Data Fig. 2. Cardiac immune cell flow analysis gating strategies.** Gating strategies for flow analysis starting with gating for immune cells ($CD45^{+}$), followed by live/dead and doublet exclusion, then gates for neutrophils ($Ly6G^{+}$), macrophages ($CD64^{+}$), CCR2 macrophages ($CCR2\text{-}GFP^{+}$), T-cells ($CD4^{+}$ or $CD8^{+}$), effector phenotype T cells ($CD4^{+}CD44^{+}$ or $CD8^{+}CD44^{+}$), dendritic cells ($Zbtb46\text{-}GFP^{+}$) or B cells ($CD19^{+}$).

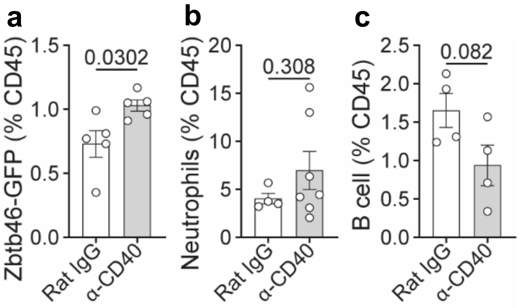


**Extended Data Fig. 3. Modest increase in dendritic cells and no significant difference in neutrophils or B cells. a,** Quantification of Zbtb46-GFP$^+$ dendritic cells in the heart by flow cytometry comparing Rat IgG (n = 5) and α-CD40 (n = 5). **b,** Quantification of neutrophils (Ly6G$^+$) in the heart by flow cytometry comparing Rat IgG (n = 4) and α-CD40 (n = 7). **c,** Quantification of B cells (CD19$^+$) in the heart by flow cytometry comparing Rat IgG (n = 4) and α-CD40 (n = 4). Two-tailed unpaired Student's *t*-test performed for all statistical analysis. Error bars indicate means ± s.e.m.

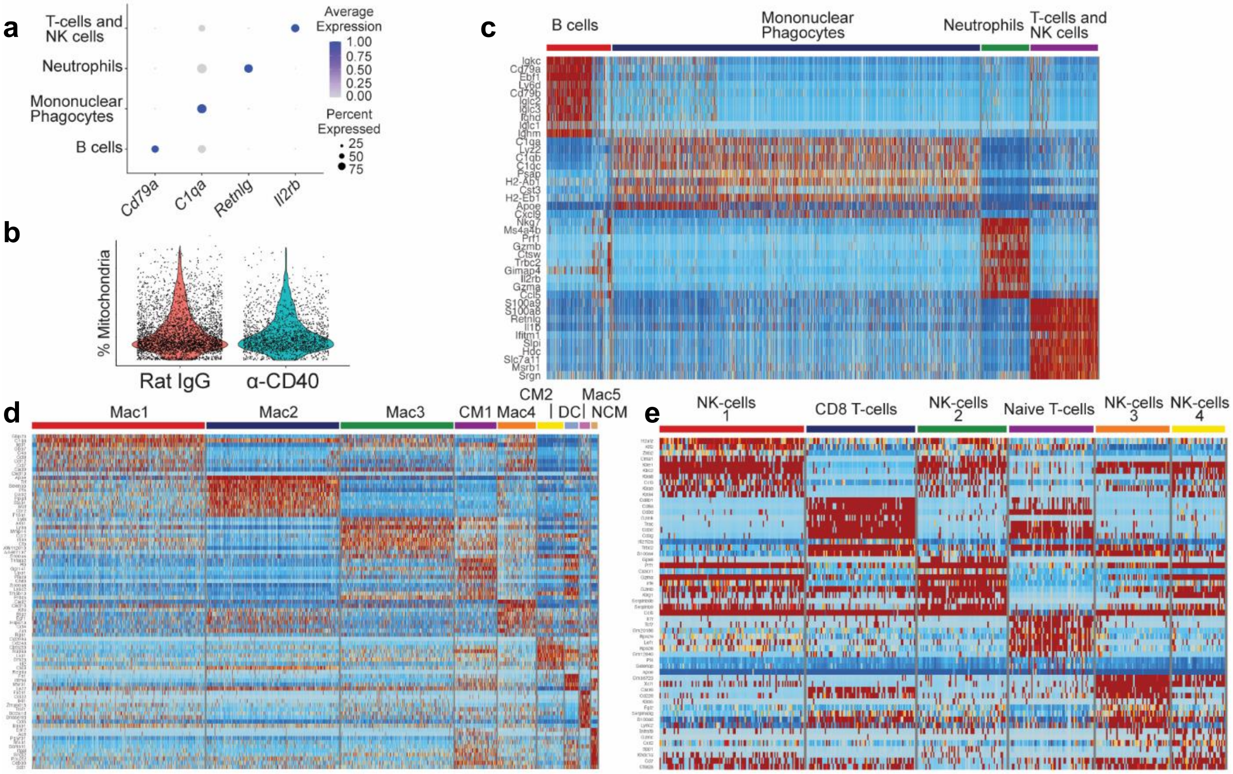


**Extended Data Fig. 4. Quality control for CITE-sequencing and gene expression signatures of major cell types and subclusters. a,** Dot plot of differentially expressed genes for the major cell types. **b,** Violin plot of percentage mitochondrial reads per cell. Cells were filtered to include those with 1,000 < RNA unique molecular identifier (UMI) count < 20,000, and mitochondrial reads less than 5%. **c,** Heatmaps showing relative expression profiles of major cell types, **d,** mononuclear phagocytes, and **e,** T-cells and NK-cells.

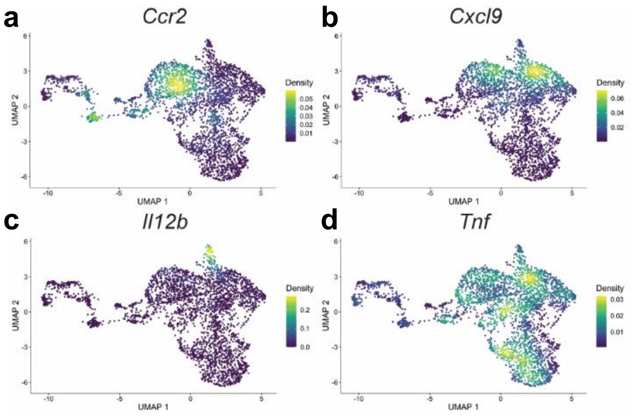


**Extended Data Fig. 5. CITE-seq expression of *Ccr2* and inflammatory cytokines and chemokines. a,** Density plot showing expression of *Ccr2*, **b,** *Cxcl9*, **c,** *Il12b*, and **d,** *Tnf* in UMAP embedding in mononuclear phagocytes from hearts treated with anti-CD40 agonist antibody.

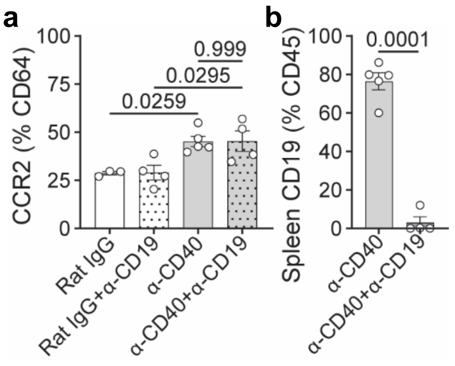


**Extended Data Fig. 6. No change in CCR2⁺ macrophage expansion with B cell depletion.**
**a,** In mice receiving depleting α-CD19 (B cell) with concurrent α-CD40 agonist or Rat IgG isotype antibodies for 7 days, $CCR2^{+}$ macrophages were quantified in the heart by flow cytometry with comparison between Rat IgG (n = 3), Rat IgG+α-CD19 (n = 4), α-CD40 (n = 5), and α-CD40+α-CD19 (n = 4). One-way ANOVA with Sidak correction. **b,** Quantification of B cells ($CD19^{+}$) in the spleen by flow cytometry confirmed B cell depletion with comparison between α-CD40 (n = 5) and α-CD40+α-CD19 (n = 4). Unpaired Student's *t*-test, two-tailed. Error bars indicate means ± s.e.m.

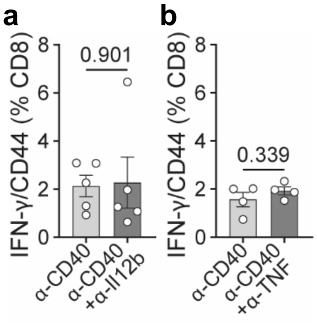


**Extended Data Fig. 7. No change in intracellular IFN-γ following CD40 agonist treatment with concurrent single agent IL12b or TNF neutralizing antibodies. a,** In mice receiving neutralizing α-IL12b with concurrent α-CD40 agonist treatment antibodies for 7 days, IFN-$\gamma^{+}$CD44$^{+}$ CD8 T-cells were quantified in the heart by flow cytometry with comparison between α-CD40 (n = 5), and α-CD40+α-IL12b (n = 5). **b,** In mice receiving neutralizing α-TNF with concurrent α-CD40 agonist treatment antibodies for 7 days, IFN-$\gamma^{+}$CD44$^{+}$ CD8 T-cells were quantified in the heart by flow cytometry with comparison between α-CD40 (n = 4), and α-CD40+α-TNF (n = 4). Two-tailed unpaired Student's *t*-test performed for all statistical analysis. Error bars indicate means ± s.e.m.

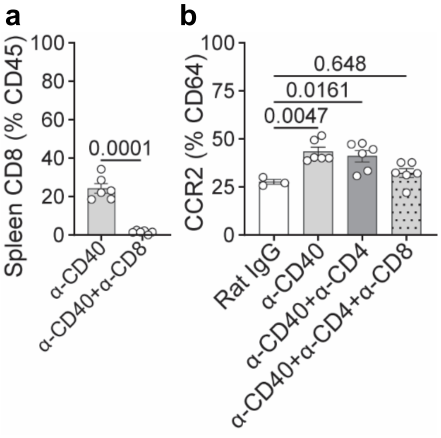


**Extended Data Fig. 8. No change in CCR2⁺ macrophage expansion with CD4 T-cell depletion. a,** In mice receiving depleting α-CD8 with concurrent α-CD40 agonist treatment antibodies for 7 days, $CD8^+$ T-cells were quantified in the spleen by flow cytometry to confirm CD8 T-cell depletion when comparing α-CD40 (n = 6) and α-CD40+α-CD8 (n = 6). Two-tailed unpaired Student's *t*-test. **b,** In mice receiving depleting single agent α-CD4 or combined α-CD4 with α-CD8 and concurrent α-CD40 agonist for 7 days, $CCR2^+$ macrophages were quantified in the heart by flow cytometry with comparison between Rat IgG (n = 3), α-CD40 (n = 6), α-CD40+α-CD4 (n = 6), and α-CD40+α-CD4+α-CD8 (n = 6). One-way ANOVA with Sidak correction. Error bars indicate means ± s.e.m.
